# Mapping of promoter usage QTL using RNA-seq data reveals their contributions to complex traits

**DOI:** 10.1101/2022.02.24.481875

**Authors:** Naoto Kubota, Mikita Suyama

## Abstract

Genomic variations are associated with gene expression levels, which are called expression quantitative trait loci (eQTL). Most eQTL may affect the total gene expression levels by regulating transcriptional activities of a specific promoter. However, the direct exploration of genomic loci associated with promoter activities using RNA-seq data has been challenging because eQTL analyses treat the total expression levels estimated by summing those of all isoforms transcribed from distinct promoters. Here we propose a computational framework for identifying genomic loci associated with promoter activities, called promoter usage quantitative trait loci (puQTL), using conventional RNA-seq data. By leveraging public RNA-seq datasets from the lymphoblastoid cell lines of 438 individuals from the GEUVADIS project, we obtained promoter activity estimates and mapped 2,592 puQTL at the 10% FDR level. The results of puQTL mapping enabled us to interpret the manner in which genomic variations regulate gene expression. We found that 310 puQTL genes (16.1%) were not detected by eQTL analysis, suggesting that our pipeline can identify novel variant–gene associations. Furthermore, we identified genomic loci associated with the activity of “hidden” promoters, which the standard eQTL studies have ignored. We found that most puQTL signals were concordant with at least one genome-wide association study (GWAS) signal, enabling novel interpretations of the molecular mechanisms of complex traits. Our results emphasize the importance of the re-analysis of public RNA-seq datasets to obtain novel insights into gene regulation by genomic variations and their contributions to complex traits.

## Introduction

Variations in the human genome generate a variety of transcriptomes across individuals. Many previous studies have identified single-nucleotide polymorphisms (SNPs) and short insertions and deletions (indels) associated with gene expression levels, which are called expression quantitative trait loci (eQTL) (Nica and Dermitzakis, 2013). eQTL are significantly enriched in genomic loci associated with complex traits identified by genome-wide association studies (GWASs), suggesting the importance of the differences in gene regulation caused by genomic variations in the context of various traits and disease risk (The GTEx Consortium, 2020). In conventional eQTL studies, the expression levels of all isoforms of each gene are quantified, and they are then merged into the total gene expression levels and used as dependent variables in linear regression analyses (Nica and Dermitzakis, 2013). If a gene has multiple promoters, it is impossible to directly assess the effects of genomic variations on the activity of each promoter using eQTL analyses. Isoforms transcribed from distinct promoters can have distinct functions even if they have the same gene name; for example *TP53* (which encodes p53) (Yang et al., 1998), *TP73* (p73) (Pozniak et al., 2000; Zaika et al., 2002), *SHC1* (p52/p46/p66) (Luzi et al., 2000; Ventura et al., 2002), *INK4a*/*ARF* (Ouelle et al., 1995; Sherr, 2001), and *CDKN1A* (p21) (Nozell and Chen, 2002). Therefore, it is essential to map the genomic variations associated with promoter activity to better understand how they affect the expression levels of specific transcripts and confer risk for complex traits.

Recently, several studies mapped the genomic variations associated with promoter activity based on cap analysis of gene expression (CAGE) technology, which captures the 5′ end cap structure of transcripts and is termed promoter usage QTL (puQTL) (Blauwendraat et al., 2016; Garieri et al., 2017) or transcription start site QTL (tssQTL) (Schor et al., 2017). Those studies have shown that puQTL or tssQTL analysis enables the detection of SNPs associated with promoter usage, which conventional eQTL analysis could not discover. However, this strategy has a high cost because it requires the generation of promoter usage data for hundreds of individuals. Thus, for mapping additional puQTL at a lower cost, it is necessary to develop a computational framework that can leverage extensive RNA-seq data from public databases.

Several RNA-seq-based studies have been performed to map transcript usage QTL (tuQTL) (Alasoo et al., 2019; Battle et al., 2013; Chen et al., 2016; Lappalainen et al., 2013; Richards et al., 2017). Most of those studies were based on transcript annotations; thus, promoters not described in standard annotation files could not be analyzed. Notably, a recent study (Alasoo et al., 2019) reported a computational method to map tuQTL by splitting data into independent events (promoter usage, splicing, and 3′ end usage), and employed Salmon (Patro et al., 2017) for quantifying full-length transcripts. Alignment-free methods, such as Salmon and kallisto (Bray et al., 2016), have been reported to have a low performance in transcript-level quantification (Srivastava et al., 2020) and to overestimate lowly expressed transcripts (Wu et al., 2018), which might lead to the mapping of false-positive puQTL. Therefore, a method that focuses on promoter activity estimates and overcomes the problems caused by the alignment-free methods is essential for accurately mapping puQTL using RNA-seq data.

This study aimed to develop a new computational framework to map puQTL by quantifying genome-wide promoter activities using publicly available RNA-seq data. In our pipeline, we employed an alignment-based method, proActiv, which has shown high performance in promoter activity estimates, as well as higher levels of agreement with H3K4me3 ChIP-seq signals compared with other methods, such as Salmon and kallisto (Demircioğlu et al., 2019). By newly constructing transcript annotations based on mapped reads, we tried to comprehensively capture the activities of promoters, including those not described in standard annotation files. Moreover, in addition to short variants (SNPs and indels), we explored associations between long variants (structural variants (SVs)) and promoter usage. We successfully mapped puQTL and found that most of them had genomic and epigenetic features that were similar to those of eQTL, in agreement with a previous report (Garieri et al., 2017). We also found that our methodology was able to detect hidden promoters associated with genomic variations, which may have been overlooked by annotation-based puQTL analyses. Some puQTL were colocalized with GWAS signals, which enabled new interpretations of complex trait associations. Overall, this study provided a successful way to identify genetic factors that regulate promoter activity by leveraging publicly available RNA-seq data. We can expand this strategy to extensive RNA-seq datasets from various tissues and cells, thus advancing our understanding of the context-specific perturbation of transcription by genomic variations.

## Methods

### Preparation of genotype and RNA-seq data

We obtained metadata and genotype data of SNPs and indels for 438 individuals from the 1000 Genomes Project (The 1000 Genomes Project Consortium, 2015) in variant call format (vcf) (hg19). Subsequently, we transformed these coordinates into hg38 coordinates using UCSC liftOver. As for SVs, we downloaded the genotype data reported by Yan et al. in vcf (hg38) (Yan et al., 2021). Then, we combined them and used the genotype data of biallelic variants with a minor allele frequency >0.01 in the samples, for downstream analysis. We obtained corresponding RNA-seq data of Epstein– Barr virus-transformed lymphoblastoid cell lines (LCLs) from the GEUVADIS project (Lappalainen et al., 2013).

### RNA-seq data processing

We performed the quality control of RNA-seq data using fastp (version 0.20.1) (Chen et al., 2018) with “-3 -q 30” options, to discard reads with low quality. We aligned the remaining reads to the human reference genome (GRCh38) and the Ensembl 104 transcript annotations using STAR (version 2.7.9a) (Dobin et al., 2013) in the two-pass mode. In the first mapping step, we used the “--outFilterMultimapNmax 1” option to allow uniquely mapped reads exclusively. In the second mapping step, we used the “--outFilterMultimapNmax 1 --sjdbFileChrStartEnd /path/to/1_SJ.out.tab/path/to/2_SJ.out.tab …. /path/to/438_SJ.out.tab --limitSjdbInsertNsj 10000000” option to use the junctions files of the first mapping as “annotated” junctions for the second mapping step. These steps allowed us to detect more spliced reads mapped to novel junctions. Using the mapped reads, we performed reference-guided transcript assembly using StringTie2 (version 2.1.7) (Kovaka et al., 2019) with the “-G /path/to/Ensembl_104_annotations.gtf --conservative” option for each sample, and merged them with the “-G /path/to/Ensembl_104_annotations.gtf --merge” option. Using the transcript annotations file and junction counts files produced by STAR, we quantified and normalized the promoter activities for each sample using proActiv (version 1.1.18) (Demircioğlu et al., 2019). We executed functions in the proActiv software in a docker container (naotokubota/proactiv:1.1.18 in Docker Hub). The promoters of single-exon genes and those that overlapped with internal exons were excluded. All promoters with promoter activity greater than zero in at least 25% of the samples were used for puQTL analysis. For the quantification of gene-level expressions, we counted reads for each transcript using featureCounts (version 1.6.4) (Liao et al., 2014) with the “-p -B -t exon -g gene_id -a /path/to/transcript_annotations_assembled_by_StringTie2.gtf” option. In accordance with the methods employed in the GTEx project (The GTEx Consortium, 2020), all genes with a transcripts per million (TPM) value >0.1 and raw read counts greater than six in at least 20% of the samples were used for eQTL analysis. The read counts of the remaining genes were normalized across samples via the trimmed mean of M values method implemented in edgeR (version 3.12.1) (Robinson et al., 2010) in a docker container (broadinstitute/gtex_eqtl:V8 in Docker Hub).

We downloaded the gene quantification data of the GM12878 cell line generated by CAGE data processing from the ENCODE project (Accession ID: ENCFF006DIB) to compare the promoter activities estimated by proActiv (Demircioğlu et al., 2019) and kallisto (Bray et al., 2016). We downloaded GM12878 RNA-seq data from the ENCODE project (Accession ID: ENCFF000EWJ, ENCFF000EWX). After quality control using fastp with the “-3 -q 30” option, we performed a random sampling of 50 million reads using SeqKit (version 0.13.2) (Shen et al., 2016) with the “sample -n 50000000 -s 20” option. We obtained the TPM value for each transcript based on the remaining reads using kallisto (version 0.44.0) and summed those of transcripts sharing the same promoter. We also obtained promoter activity scores using proActiv, as well as GEUVADIS RNA-seq data processing. We excluded promoters with zero activity in all methods (CAGE, kallisto, and proActiv) and calculated Spearman’s ρ for each pair of promoter activity sets.

### QTL mapping

We calculated probabilistic estimation of expression residuals (PEER) factors (Stegle et al., 2012) based on the promoter activity and gene expression matrix for the puQTL and eQTL analysis using the PEER package (version 1.0) in a docker container (broadinstitute/gtex_eqtl:V8 in Docker Hub). For QTL discovery, we used QTLtools (Delaneau et al., 2017) (version 1.3.1) in a docker container (naotokubota/qtltools:1.3.1 in Docker Hub). We calculated genetic principal components (PCs) from the genotype matrix using QTLtools pca with the “--scale --center --maf 0.01 --distance 50000” option. First, we used the nominal pass using QTLtools cis with the “--nominal 0.00001 --normal” option, to check the suitable number of PEER factors (as covariates) that were necessary to improve sensitivity. We checked the number of QTL with nominal *P*-values <1.0 · 10^−5^ using varying numbers of PEER factors (0, 1, 2, 3, 4, 5, 6, 7, 8, 9, 10, 15, 20, 25, 30, 35, 40, 45, and 50). Next, we performed a permutation pass using QTLtools cis with the “--permute 1000 --normal --std-err” option. We included the top five genetic PCs, sex, and the suitable number of PEER factors for each category (25 for puQTL and 50 for eQTL) as covariates in the QTL analysis. Subsequently, using the output of the permutation pass, we performed a false discovery rate (FDR) correction on the permutation *P*-values at the 10% FDR level using “Rscript /qtltools/scripts/qtltools_runFDR_cis.R.” Finally, we performed a conditional pass to obtain the significant QTL at the 10% FDR level using QTLtools cis with the “--mapping /path/to/thresholds.txt --normal --std-err” option. We also performed a nominal pass using QTLtools cis with the “--nominal 1.0 --normal --std-err” option, to obtain the statistics of all associations in a ±1 Mb *cis* window. To estimate causal variants for puQTL, we performed fine-mapping using SuSiE (version 0.11.42) (Wang et al., 2020; Zou et al., 2021) in a docker container (naotokubota/coloc-locuscomparer:1.0 in Docker Hub). We obtained the posterior inclusion probability (PIP) for each variant–phenotype pair using the “susie_rss()” function. We used the UCSC Genome Browser (Haeussler et al., 2019) to visualize the genomic positions around the QTL.

### HiChIP data processing

We downloaded the published data of H3K27ac HiChIP of the GM12878 cell line (GEO accession number: GSM2705041) (Mumbach et al., 2017) and performed quality control using fastp (version 0.20.1) (S. Chen et al., 2018) with the “-3 -q 30” option, to discard reads with low quality, as well as RNA-seq data processing. We aligned the remaining reads to the human reference genome (GRCh38) using the HiC-Pro pipeline (version 3.0.0) (Servant et al., 2015) in a docker container (nservant/hicpro:3.0.0 in Docker Hub). We used the default settings to remove duplicate reads, assign reads to MboI restriction fragments, and filter valid interactions. We performed peak calling using MACS2 (Zhang et al., 2008) with the “callpeak -f BAM -g hs -q 0.01” option and obtained sets of high-confidence loops (5 kb bin, 1% FDR level) using FitHiChIP (version 7.0) (Bhattacharyya et al., 2019) with the configfile_BiasCorrection_CoverageBias file and default settings in a docker container (aylab/fithichip:latest in Docker Hub).

### Genomic and epigenetic enrichment analysis

We downloaded the peak and signal files of ChIP-seq of histone marks and ATAC-seq of the GM12878 cell line from the ENCODE project (Dunham et al., 2012) (Accession ID: H3K4me3, ENCFF587DVA; H3K4me1, ENCFF321BVG; H3K27ac, ENCFF023LTU; H3K9ac, ENCFF069KAG; H3K27me3, ENCFF291DHI; ATAC-seq, ENCFF748UZH), transcription factor (TF) footprinting data based on DNase-seq of the GM12865 and GM06990 cell lines (Vierstra et al., 2020), and the chromatin states data of the GM12878 cell line defined by ChromHMM (Ernst and Kellis, 2017) from the Roadmap Epigenomics Project (Roadmap Epigenomics Consortium et al., 2015). The chromatin states fell into 25 categories: 1_TssA (Active TSS), 2_PromU (Promoter Upstream TSS), 3_PromD1 (Promoter Downstream TSS 1), 4_PromD2 (Promoter Downstream TSS 2), 5_Tx5’’ (Transcribed - 5’ preferential) 6_Tx (Strong transcription), 7_Tx3’’ (Transcribed - 3’ preferential), 8_TxWk (Weak transcription), 9_TxReg (Transcribed & regulatory (Prom/Enh)), 10_TxEnh5’’ (Transcribed 5’ preferential and Enh), 11_TxEnh3’’ (Transcribed 3’ preferential and Enh), 12_TxEnhW (Transcribed and Weak Enhancer), 13_EnhA1 (Active Enhancer 1), 14_EnhA2 (Active Enhancer 2), 15_EnhAF (Active Enhancer Flank), 16_EnhW1 (Weak Enhancer 1), 17_EnhW2 (Weak Enhancer 2), 18_EnhAc (Primary H3K27ac possible Enhancer), 19_DNase (Primary DNase), 20_ZNF/Rpts (ZNF genes & repeats), 21_Het (Heterochromatin), 22_PromP (Poised Promoter), 23_PromBiv (Bivalent Promoter), 24_ReprPC (Repressed Polycomb), and 25_Quies (Quiescent/Low). Moreover, we tested whether a set of QTL is significantly enriched in the functional annotations using QTLtools fenrich with the “--permute 1000” option. We used the positions of best hit variants for each QTL and those of all promoters (*n* = 12,957) and all genes (*n* = 12,275) for the QTLtools fenrich analysis. We generated aggregation plots of the ChIP-seq and ATAC-seq profiles of the GM12878 cell line (Accession ID: H3K4me3, ENCFF927KAJ; H3K4me1, ENCFF564KBE; H3K27ac, ENCFF469WVA; H3K9ac, ENCFF028KBY; H3K27me3, ENCFF919DOR; ATAC-seq, ENCFF603BJO) using deepTools (version 3.5.1) (Ramírez et al., 2016) in a docker container (quay.io/biocontainers/deeptools:3.5.1--py_0 in Quay.io) with the positional information of the significant QTL in the bed format.

### GWAS hit enrichment and colocalization analysis

We tested whether the QTL variants are concordant with GWAS lead variants using QTLtools rtc with the “--hotspots /path/to/hotspots.bed --normal --gwas-cis” option. We downloaded the recombination hotspot data from the 1000 Genomes Project and GWAS lead variant data from the GWAS Catalog (MacArthur et al., 2017) for the QTLtools rtc analysis. We also performed a colocalization analysis using coloc (version 5.1.0) (Wallace, 2021) in a docker container (naotokubota/coloc-locuscomparer:1.0 in Docker Hub) with the GWAS summary statistics obtained from the GWAS Catalog. We calculated *r*-squared values (*r*^2^) between variants of interest and variants around them using PLINK (version 1.90b6.21) (Purcell et al., 2007) with the “--r2 --ld-window-kb 1000 --ld-window 99999 --ld-window-r2 0” option in a docker container (quay.io/biocontainers/plink:1.90b6.21--h779adbc_1 in a Quay.io).

### Comparison of multiple protein sequences

We downloaded the protein sequences of interest from UniProt (https://www.uniprot.org) and performed multiple alignments of protein sequences using MUSCLE (https://www.ebi.ac.uk/Tools/msa/muscle/) (Edgar, 2004). We searched for protein domains in the sequences using InterPro (https://www.ebi.ac.uk/interpro/) (Blum et al., 2020). To predict protein structures from the sequences, we used ColabFold (Mirdita et al., 2021), which is a free platform of AlphaFold2 coupled with Google Colaboratory.

### Computational environments

The computational framework was developed in the GNU Bash 3.2 environment. We generated all graphs using the pandas (version 1.1.3), matplotlib (version 3.3.1), and seaborn (version 0.11.0) packages in the Python environment (version 3.8.5). Furthermore, we used the scipy (version 1.6.2) package for all statistical tests, and docker (version 20.10.7) for the stable execution of various software.

## Results

### Systematic identification of puQTL using RNA-seq data

We aimed to identify genomic variations associated with promoter activities (puQTL) and total gene expression levels (eQTL) using RNA-seq data (Figure 1A). First, we checked whether it is possible to estimate promoter activity accurately from RNA-seq data. We compared the promoter activity of the GM12878 cell line measured by CAGE with those estimated by proActiv (Demircioğlu et al., 2019) and kallisto (Bray et al., 2016), which can accept RNA-seq data as input. Our results also showed that the promoter activity estimated by proActiv was correlated more strongly with that measured by CAGE vs. kallisto (Spearman’s ρ = 0.647 and 0.432 for proActiv and kallisto, respectively) (Figure S1). The proActiv software has been reported to exhibit high performance in promoter usage inference compared with other methods, and to capture changes in alternative promoter usage in cancer transcriptomes (Demircioğlu et al., 2019). Although proActiv is limited in that it cannot estimate the activities of promoters that overlap with internal exons, we obtained a considerable advantage in accuracy and confirmed that it is possible to utilize publicly available RNA-seq data from various resources; thus, we employed proActiv for the estimation of promoter activity in this study. Next, we developed computational pipelines for puQTL and eQTL identification using the RNA-seq dataset of LCLs from 438 individuals provided by the GEUVADIS project (Lappalainen et al., 2013) (Figure 1B, S2; Table S1). After quality control, we aligned the RNA-seq reads to the human reference genome in the two-pass mode implemented in STAR to obtain as many junction reads as possible, because the estimation of promoter activity by proActiv depends on the number of junction reads mapped on the first and second exon. Using the mapped reads of 438 individuals and the Ensembl gene annotation file, we re-constructed gene annotations using StringTie2 as an input file of proActiv. These steps allowed us to gain additional promoters to be analyzed, because proActiv can only detect the promoters described in the input GTF file. Using datasets of normalized promoter activities, gene expression levels, and genotypes, we mapped puQTL and eQTL using QTLtools. To maximize the number of significant puQTL signals, we used the top five genetic PCs, sex, and PEER factors (Stegle et al., 2012) calculated based on phenotype tables as covariates in the regression analysis (Figure S3). To conduct a comprehensive survey of the genetic factors associated with gene regulation, here we used genotypes of not only short variants, such as SNPs and indels, but also SVs recently discovered by combined analysis of short-read and long-read sequencing data (Yan et al., 2021). As a result, we successfully mapped 2,592 puQTL and 18,205 eQTL at the 10% FDR level. We observed substantial deviations of the resulting *P*-values from the null expectation for both puQTL and eQTL, indicating that our analysis was well calibrated (Figure 1C, S4A). We found that puQTL and eQTL were likely to be located near target promoters (Figure 1D, S4B), in concordance with previous reports (Garieri et al., 2017). Moreover, although we encountered a limitation in that internal promoters were excluded from our analysis, we confirmed that our results were consistent with those of the CAGE-based puQTL analysis (Garieri et al., 2017); i.e., an intergenic variant, rs8028374 (A/G), was mapped as puQTL associated with the most external promoters of the *TTC23* gene, prmtr.35339 (*P* = 6.99 · 10^−18^, β = −0.56) (Figure S5A, B), and an exonic variant, rs35430374 (C/A), was not mapped as puQTL associated with external promoters of the *DENND2D* gene (*P* = 7.78 · 10^−5^, β = −0.41 for prmtr.53803; *P* = 0.22, β = −0.13 for prmtr.53804) (Figure S5C, D).

**Figure 1.**
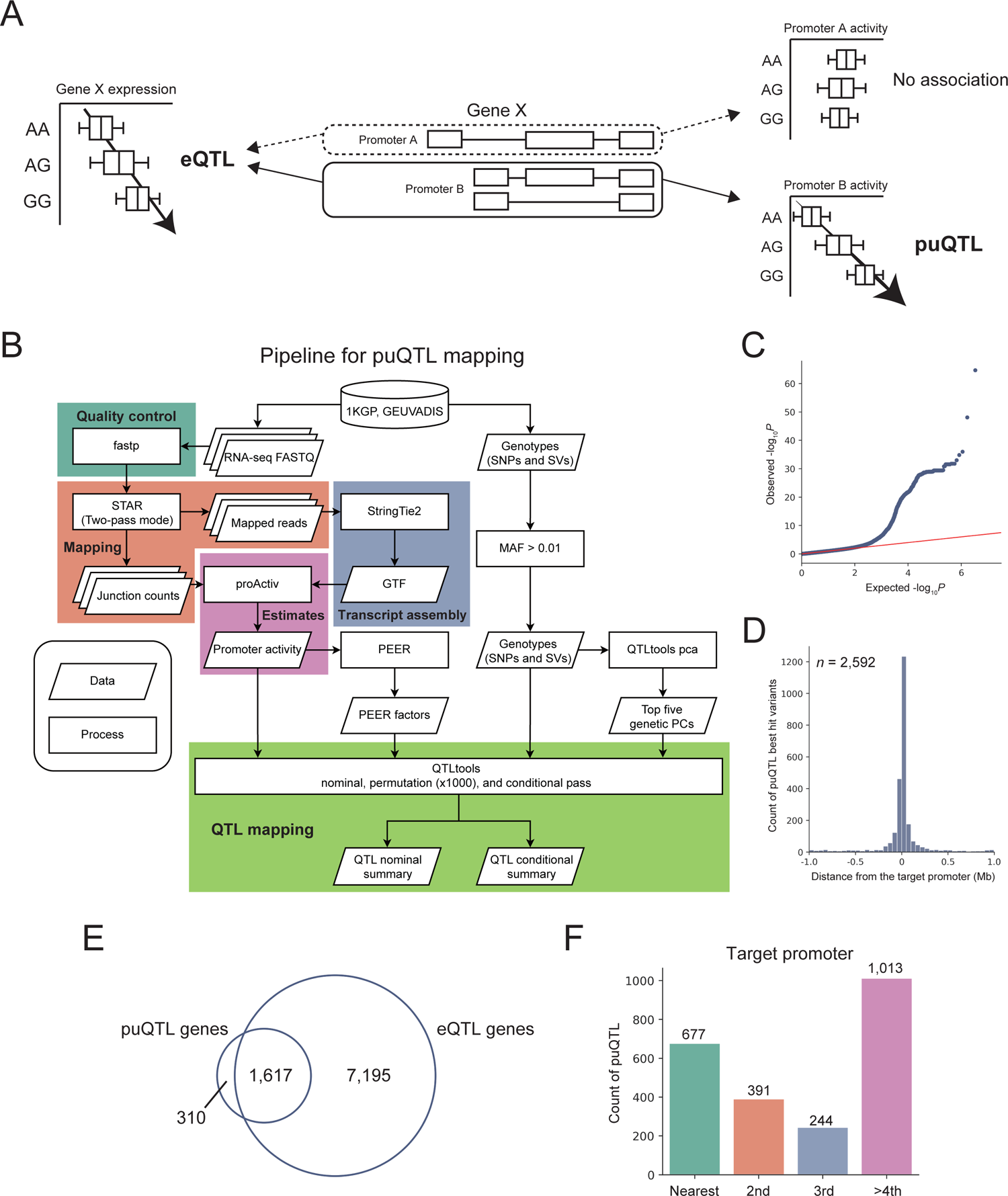
puQTL mapping using RNA-seq data. (A) Schematic view of eQTL and puQTL mapping. (B) Pipeline for puQTL mapping. 1KGP, the 1000 Genomes Project; MAF, minor allele frequency; GTF, gene transfer format. (C) Quantile–quantile plot of *P*-values. The nominal pass results of chromosome 22 are plotted, and the red line indicates the expected *P*-values under the null hypothesis. (D) Distribution of the distance of puQTL best hit variants from the target promoters. (E) Overlap of puQTL genes and eQTL genes. (F) Counts of puQTL associated with multi-promoter genes based on categories of proximity to their target promoters. “Nearest” means that the puQTL targets the nearest promoters of the associated genes, “2nd” means that they target the second nearest promoters, and so forth.

Although the number of puQTL was smaller than those of the previous CAGE-based puQTL analysis because of the limited power of RNA-seq-based estimates of promoter activities, these results demonstrated that our pipeline provides promising results. We found that 16.1% of the genes associated with puQTL (puQTL genes) (310/1,927) did not overlap with those associated with eQTL (eQTL genes) (Figure 1E), suggesting that puQTL mapping enables the identification of overlooked variant–gene associations. We also performed fine-mapping of puQTL to estimate candidate causal variants using the SuSiE software (Wang et al., 2020; Zou et al., 2021). We identified 424 variants with a high causality (PIP > 0.9) and found that two of the credible causal variants were SVs (967_HG00733_ins and 28764_HG02059_ins), suggesting their independent roles in the perturbation of promoter usage.

Unlike eQTL, the puQTL analysis has the advantage of predicting associations between promoters and genomic variations. We expected that puQTL associated with multi-promoter genes would target the nearest promoter. However, the proportion of puQTL targeting the nearest promoters in all puQTL associated with multi-promoter genes was 29.1% (677/2,325) (Figure 1F). This result suggests that the effects of genetic factors on promoter usage are not easy to dissect by simply focusing on genomic distances, which emphasizes the importance of puQTL mapping.

### puQTL are enriched in active promoters and enhancers

Next, we sought to examine the epigenetic contexts of puQTL to understand their functional mechanisms in gene regulation. We found that active histone marks for transcription (H3K4me3, H3K27ac, and H3K9ac) and ATAC-seq signals, which represent the open chromatin structure, were concentrated around puQTL, as well as eQTL (Figure 2A). We assessed the statistical significance of the enrichment in epigenetic features based on permutation tests using the QTLtools fenrich module. The results of this analysis showed that puQTL and eQTL were significantly enriched in active histone marks, open chromatin regions detected by ATAC-seq, and TF footprints (Figure 2B, C). We confirmed that puQTL that were discovered without PEER factors as covariates were not enriched in transcriptionally active regions (Figure S6), whereas those discovered using 25 PEER factors were enriched in those regions (Figure 2B). This result suggests that, although the greatest number of puQTL was detected when we used no PEER factors (Figure S3B), the puQTL discovered using 25 PEER factors were more credible than those identified without PEER factors. We also tested if the sets of QTL are enriched in regulatory elements using the chromatin states data of the GM12878 cell line defined by ChromHMM (Ernst and Kellis, 2017). We found that both puQTL and eQTL were significantly enriched in upstream promoter regions (2_PromU) and active enhancer elements (13_EnhA1) at the 5% FDR level, implying the functional roles of the genomic variations in gene regulation via *cis*-regulatory elements, such as promoters and enhancers (Figure 2D, E). These results showed that the general features of chromatin states are similar between puQTL and eQTL.

**Figure 2.**
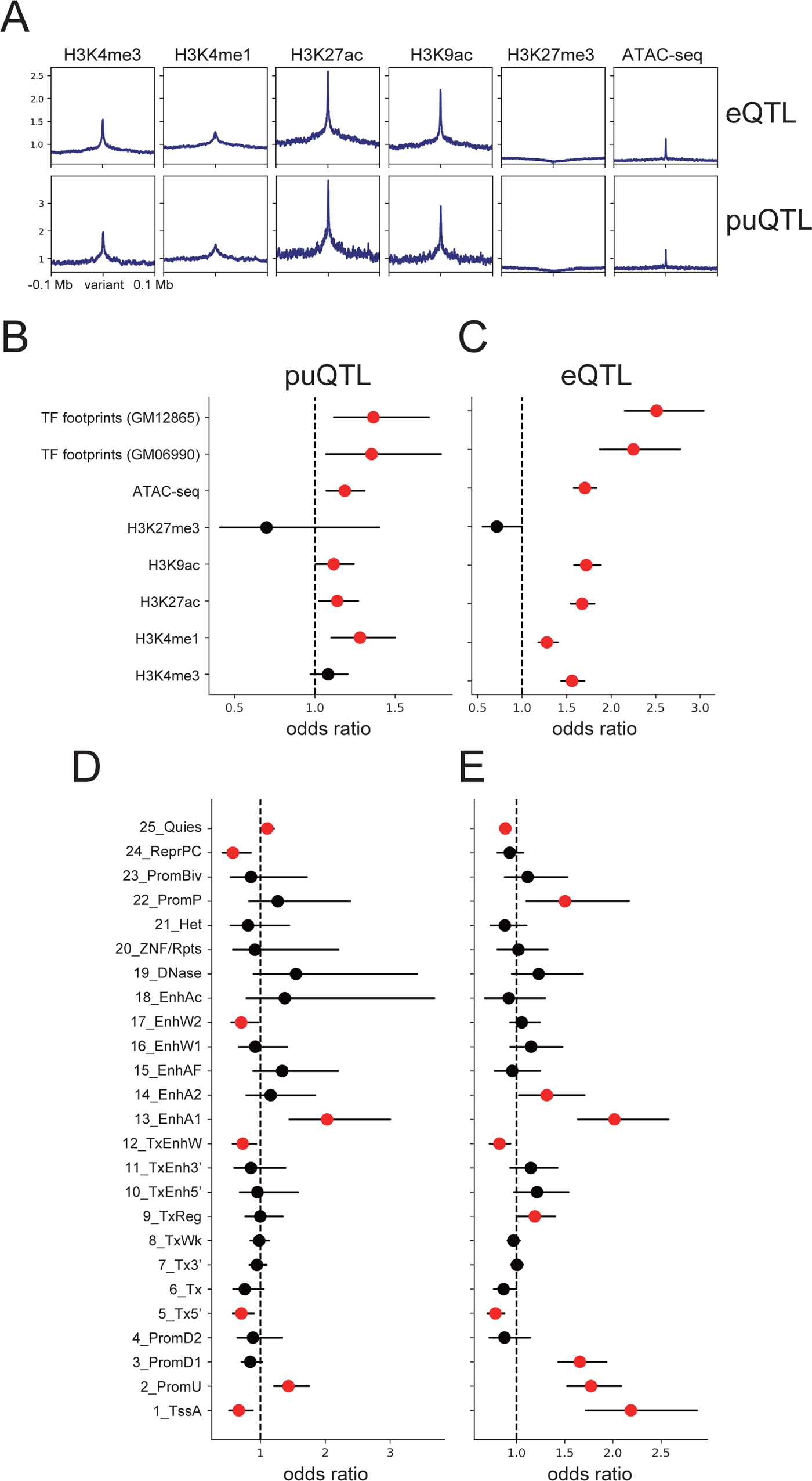
Enrichment of epigenetic features. (A) Aggregation plots of histone mark ChIP-seq and ATAC-seq signals for the 200 kb regions flanking puQTL best hit variants. (B–E) Enrichment of the peaks of histone mark ChIP-seq and transcription factor footprints at puQTL (B) and eQTL (C), and enrichment of the chromatin states defined by ChromHMM (Ernst and Kellis, 2017) at puQTL (D) and eQTL (E). The red dots represent significant enrichment at the 5% FDR level, and the bars indicate 95% confidence intervals.

### puQTL-associated gene classification

Next, we classified genes associated with puQTL into four groups in a similar way as that described by a previous study (Garieri et al., 2017) (Figure 3A). Group-1 included 212 genes with a single promoter. Among the multi-promoter genes, 1,626 genes harboring only one promoter associated with puQTL were classified into group-2. Considering effect direction, which is a slope in the linear regression analysis (β), group-3 included 32 genes associated with puQTL with opposite effects on distinct promoters, and group-4 included 57 genes associated with puQTL with concordant effects on distinct promoters (Figure 3B). Based on this classification, we unveiled the detailed mechanisms of gene regulation by genomic variations.

**Figure 3.**
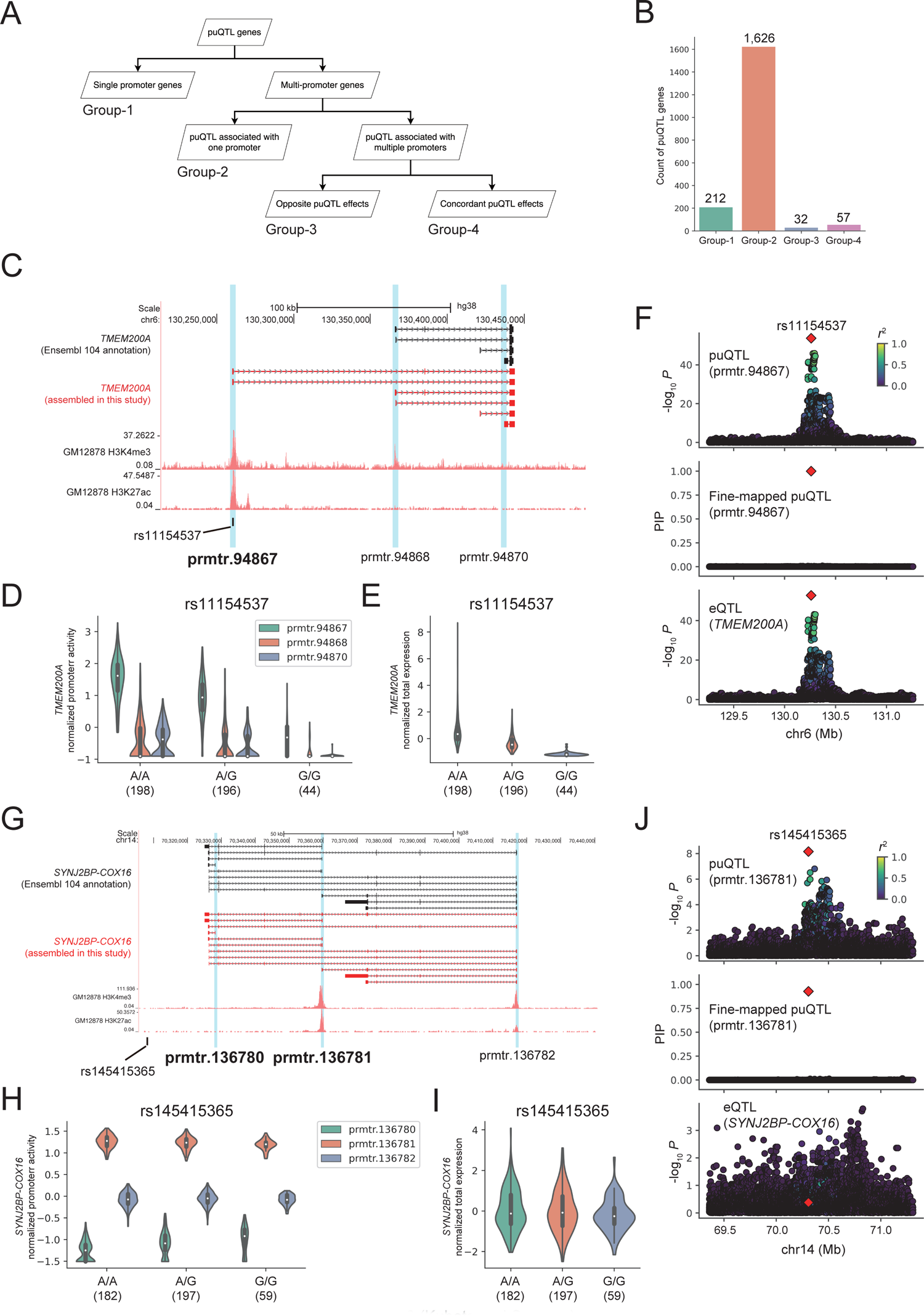
puQTL-associated gene classification. (A) The procedure used for puQTL gene classification. (B) Count of each group of puQTL genes. (C) The *TMEM200A* gene locus. The structures of the *TMEM200A* gene in the Ensembl 104 annotation and assembled in this study are presented in black and red, respectively, with ENCODE GM12878 H3K4me3 and H3K27ac ChIP-seq signals. The vertical blue bars indicate the location of active promoters. The black bar indicates the location of the rs11154537 variant. (D, E) Comparison of the promoter activities (D) and total expression levels (E) of the *TMEM200A* gene among the rs11154537 genotypes. The numbers in parentheses indicate sample size. (F) The associations of puQTL, fine-mapped puQTL for prmtr.94867, and eQTL for *TMEM200A* are shown in the top, middle, and bottom panels. rs11154537 is plotted in a red diamond, and the colors indicate the *r*-squared values between rs11154537 and other variants. (G) The *SYNJ2BP-COX16* gene locus. (H, I) The promoter activities (H) and total expression levels (I) of the *SYNJ2BP-COX16* gene were compared among the rs145415365 genotypes. (J) Associations of puQTL, fine-mapped puQTL for prmtr.136781, and eQTL for *SYNJ2BP-COX16* are shown in the top, middle, and bottom panels.

We found an illustrative puQTL that was overlooked by annotation-based QTL analyses. Among the puQTL genes in group-4, the *TMEM200A* gene (Figure 3C) encodes a member of the transmembrane (TMEM) protein family. This gene has three active promoters (prmtr.94867, prmtr.94868, and prmtr.94870). Notably, isoforms transcribed from prmtr.94867 were not described in the Ensembl gene annotation file and were newly added by transcript assembly in our pipeline; therefore, the previous annotation-based analysis could not detect this promoter. The H3K4me3 and H3K27ac ChIP-seq signals also supported the contention that prmtr.94867 is transcriptionally active in the GM12878 cell line (Figure 3C). We found that rs11154537 (A/G), which is a variant located in the first exon of the isoform transcribed by prmtr.94867, was mapped as puQTL associated with the activity of prmtr.94867 (*P* = 1.96 · 10^−54^, β = −0.99) and prmtr.94870 (*P* = 8.88 · 10^−8^, β = −0.38), but not with prmtr.94868 (*P* = 5.40 · 10^−4^, β = −0.25) (Figure 3D). This variant was also mapped as eQTL associated with the total expression level of the *TMEM200A* gene (*P* = 1.93 · 10^−53^, β = −0.99) (Figure 3E). In addition, the result of the puQTL fine-mapping showed that the PIP of rs11154537, which represents the estimated causality in the perturbation of the promoter activity, was 1.00 for prmtr.94867 (Figure 3F), strongly suggesting that this variant is causal of the perturbation of prmtr.94867 usage. Taken together, these results indicate that an alternative allele of rs11154537 somehow leads to the downregulation of the “hidden” major promoter activity (prmtr.94867), resulting in a decrease in the total expression level of the *TMEM200A* mRNA. This is an example of the successful interpretation of how genomic variations alter total gene expression levels. Moreover, this example supports the conclusion that our computational pipeline is sufficiently powerful for detecting hidden active promoters associated with genomic variations, which were not included in eQTL and puQTL analyses based on standard annotation files.

Two puQTL genes associated with fine-mapped SVs were included in group-2. One of them is the *IVNS1ABP* gene, which encodes the influenza virus NS1A-binding protein. This gene was actively transcribed from one promoter, prmtr.86427 (Figure S7A). We found that 967_HG00733_ins, a genomic insertion of 34 bp, was located in the 90 kb upstream region of the gene and mapped as puQTL associated with the activity of prmtr.86427 (*P* = 1.71 · 10^−12^, β = 0.48) (Figure S7B). This SV was also mapped as eQTL associated with the total expression level of the *IVNS1ABP* gene (*P* = 4.61 · 10^−56^, β = 0.96) (Figure S7C). The chromatin interactions detected using H3K27ac HiChIP data of the GM12878 cell line (Mumbach et al., 2017) also support the functional connections between the SV and prmtr.86427 (Figure S7A). The PIP of 967_HG00733_ins was 0.99 for prmtr.86427 (Figure S7D). These results demonstrated that the intergenic 34 bp insertion upregulates the activity of one promoter via a 90 kb-range enhancer, thus increasing the total expression level of the *IVNS1ABP* mRNA. *RSPH1* was another puQTL gene associated with a fine-mapped SV. This gene encodes a radial-spoke-head protein and is transcribed from two distinct promoters (prmtr.70068 and prmtr.70069) (Figure S7E). Notably, the isoforms transcribed from prmtr.70068 were newly detected by transcript assembly in our pipeline, as in the case of the *TMEM200A* gene (Figure 3C). A 166 bp insertion located within the first exon of the isoform transcribed from prmtr.70068, 28764_HG02059_ins, was mapped as puQTL associated with the activity of prmtr.70068 (*P* = 2.75 ·10^−45^, β = −0.92), but not with prmtr.70069 (*P* = 0.091, β = −0.12) (Figure S7F). This SV was also mapped as eQTL associated with the total expression level of the *RSPH1* gene (*P* = 6.77 · 10^−44^, β = −0.91) (Figure S7G). Moreover, the PIP of 28764_HG02059_ins was 1.00 for prmtr.70068 (Figure S7H), strongly suggesting that this exonic SV somehow leads to the downregulation of the promoter activity and a decrease in total *RSPH1* expression. These examples showed that our pipeline could identify cases in which SVs drive the dysregulation of promoter activities.

We also found several interesting examples of promoter usage changes without a gene expression change in group-3. One example was associated with the *SYNJ2BP-COX16* gene, which harbors three active promoters (prmtr.136780, prmtr.136781, and prmtr.136782) (Figure 3G). rs145415365 (A/G), a variant located in the downstream intergenic region of this gene, was mapped as puQTL associated with the activity of two promoters with opposite effects (*P* = 1.67 · 10^−10^, β = 0.44 for prmtr.136780; *P* = 7.02 · 10^−9^, β = −0.40 for prmtr.136781), but not with another promoter (*P* = 0.52, β = 0.045 for prmtr.136782) (Figure 3H). Notably, this variant was not associated with the total expression level of the *SYNJ2BP-COX16* gene (*P* = 0.21, β = −0.087) (Figure 3I), and the results of fine-mapping revealed a PIP of 0.93 for prmtr.136781 (Figure 3J), indicating the causality of this variant in the perturbation of promoter usage. We found another example of puQTL that was not mapped as eQTL in the *MUC12-AS1* gene. This gene harbors three active promoters (prmtr.99696, prmtr.99698, and prmtr.99699), two of which (prmtr.99696 and prmtr.99699) were not described in the Ensembl gene annotation file and were newly detected by our computational pipeline (Figure S8A). rs10229453 (T/C), a variant located in the intronic region of this gene, was mapped as puQTL associated with all promoters, but their effect directions were opposite (*P* = 2.25 · 10^−56^, β = −0.95 for prmtr.99696; *P* = 5.02 · 10^−20^, β = 0.60 for prmtr.99698; *P* = 4.02 · 10^−9^, β = 0.40 for prmtr.99699) (Figure S8B). This variant was not associated with the total expression level of the *MUC12-AS1* gene (*P* = 0.21, β = −0.087) (Figure S8C). These associations had not been detected by standard eQTL studies, which emphasizes the importance of our puQTL analysis for the discovery of new variant– gene associations.

### puQTL analyses enable novel interpretations of GWAS associations

To assess the involvement of puQTL in complex traits, we performed a comparative analysis of puQTL and GWAS lead variants. First, we applied the regulatory trait concordance (RTC) (Nica et al., 2010) method implemented in QTLtools to the set of puQTL and all GWAS lead variants curated by the GWAS Catalog. We found 1,690 puQTL, including 61 variants not mapped as eQTL, with a high-confidence concordance threshold (RTC > 0.9) for at least one trait. The set of puQTL included 166 credible causal variants (PIP > 0.9), implying the functional connections between promoter usage and traits. We found that a fine-mapped variant, rs2382817 (A/C) (PIP = 0.96), can be causal regarding the risk of inflammatory bowel diseases (IBDs), including ulcerative colitis and Crohn’s disease. Previous studies have reported that the alternative allele of rs2382817 (rs2382817-C) is protective against IBD risk (Liu et al., 2015; Jostins et al., 2012; Lange et al., 2017). The variant was mapped as puQTL associated with two active promoters of the *TMBIM1* gene (*P* = 3.50 · 10^−68^, β = 1.03 for prmtr.64998; *P* = 1.71 · 10^−9^, β = 0.41 for prmtr.65000) (Figure 4A, B), and was also mapped as eQTL associated with the total gene expression (*P* = 1.62 · 10^−41^, β = 0.85) (Figure 4C). In addition, we performed the colocalization analysis implemented in the coloc software (Wallace, 2021) to test if a causal variant can drive both puQTL and GWAS association. We found that a GWAS association for IBD (Liu et al., 2015) showed a high colocalization probability with the puQTL association for prmtr.64998 (PP4 = 0.90) and with the eQTL association for the *TMBIM1* gene (PP4 = 0.90) (Figure 4D). Taken together, these results demonstrate that an alternative allele of rs2382817 (rs2382817-C) might upregulate the minor promoter activity (prmtr.64998) and lead to a total increase in *TMBIM1* gene expression, resulting in a reduction of IBD risk. The TMBIM1 protein, which is also known as RECS1, is located in membranous compartments, including lysosomes, endosomes (Zhao et al., 2006), and the Golgi apparatus (Lisak et al., 2015), and plays a protective role in Fas-mediated apoptosis by reducing Fas expression at the plasma membrane (Shukla et al., 2011). This is an illustrative example that we can clearly interpret the perturbed promoter in the context of disease risk.

**Figure 4.**
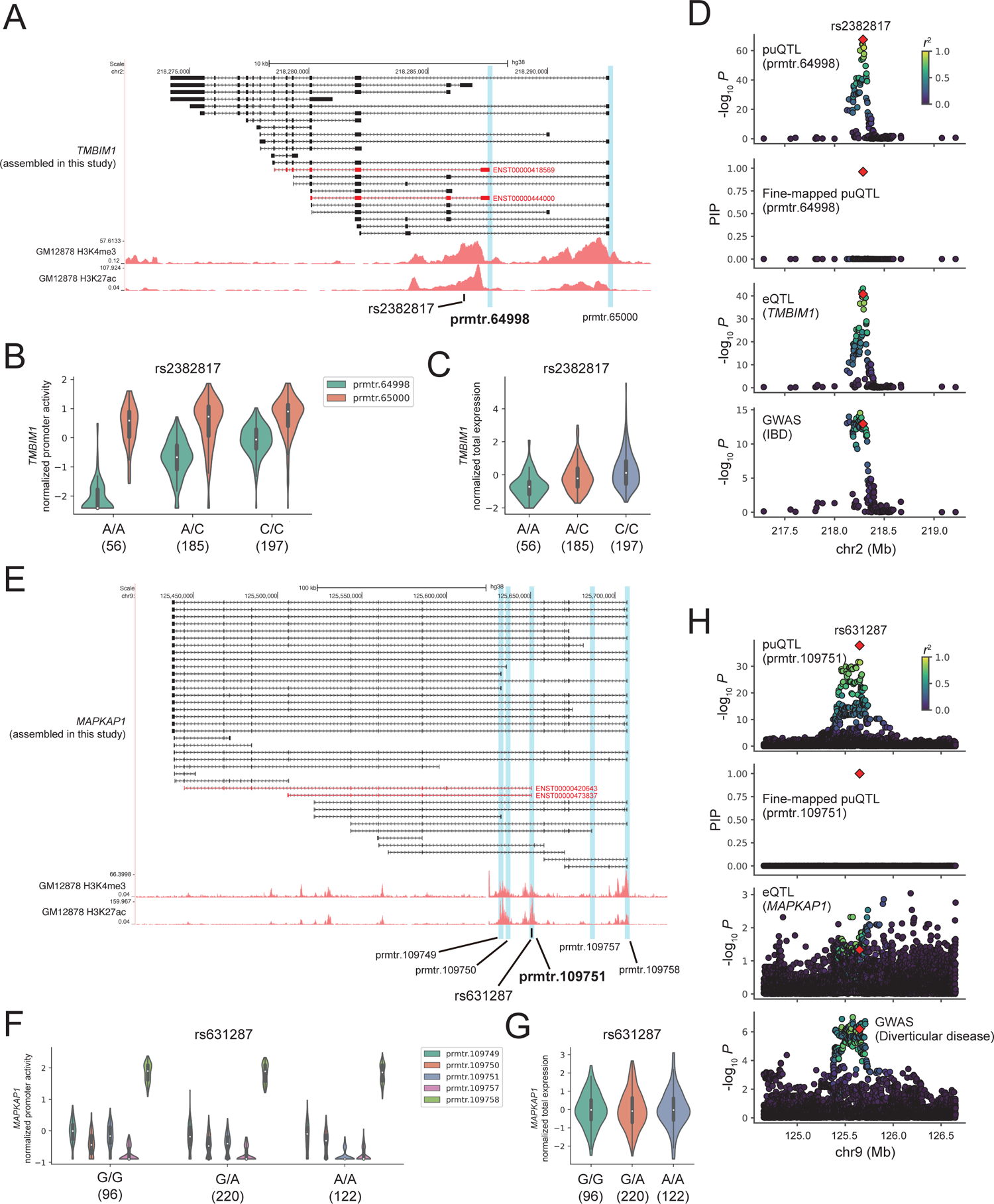
puQTL and GWAS associations. (A) The *TMBIM1* gene locus. Structures of the *TMBIM1* assembled in this study are presented in black with ENCODE GM12878 H3K4me3 and H3K27ac ChIP-seq signals. The isoforms transcribed from prmtr.64998 are shown in red. The vertical blue bars indicate the location of active promoters. The black bar indicates the location of the rs2382817 variant. (B, C) Comparison of the promoter activities (B) and total expression levels (C) of the *TMBIM1* gene among the rs2382817 genotypes. The numbers in parentheses indicate sample size. (D) The associations of puQTL, fine-mapped puQTL for prmtr.64998, eQTL for *TMBIM1*, and GWAS of inflammatory bowel disease (IBD) are shown from the top to bottom panels. rs2382817 is plotted in a red diamond, and the colors indicate *r*-squared values between rs2382817 and other variants. (E) The *MAPKAP1* gene locus. The structures of the *MAPKAP1* gene assembled in this study are presented in black, and the isoforms transcribed from prmtr.109751 are shown in red. (F, G) Comparison of the promoter activities (F) and total expression levels (G) of the *MAPKAP1* gene among the rs631287 genotypes. (H) The associations of puQTL, fine-mapped puQTL for prmtr.109751, eQTL for *MAPKAP1*, and GWAS of diverticular disease are shown from the top to bottom panels.

Importantly, our pipeline was able to demonstrate that puQTL enables novel findings of variant–gene–trait associations. We found that the rs631287 (G/A) variant mapped as puQTL associated with one promoter (prmtr.109751) of the *MAPKAP1* gene (*P* = 1.64 · 10^−38^, β = −0.80), but not associated with other promoters (*P* = 0.012, β = −0.17 for prmtr.109748; *P* = 0.54, β = 0.042 for prmtr.109750; *P* = 0.57, β = −0.039 for prmtr.109757; and *P* = 0.015, β = 0.16 for prmtr.109758) (Figure 4E, F). The prmtr.109751 is one of the minor promoters of the *MAPKAP1* gene and does not account for a large portion of the total expression; therefore, rs631287 was not mapped as eQTL for the gene (*P* = 0.046, β = 0.14) (Figure 4G). We found that a GWAS association of diverticular disease (Maguire et al., 2018), which is a condition that involves small pockets called diverticula that form from the wall of the colon, showed high colocalization probability with the puQTL association for prmtr.109751 (PP4 = 0.90); however, we observed no colocalization with the eQTL association for the *MAPKAP1* gene (PP4 = 0.091) (Figure 4H), indicating that standard eQTL studies cannot detect this variant–gene–trait association. The *MAPKAP1* gene, also known as *SIN1*, encodes the evolutionarily conserved protein Sin1, which is an essential component of the mammalian TOR complex 2 (mTORC2) (Jacinto et al., 2006; Yang et al., 2006). The full-length *MAPKAP1* mRNA is translated into a protein that contains three domains; i.e., N-terminal domain (NTD), conserved region in middle domain, and Pleckstrin-homology domain (SIN1_HUMAN) (Figure S9A, B). However, we found that one isoform transcribed by prmtr.109751 (ENST00000420643) was predicted to be translated into an N-terminally truncated protein lacking the NTD sequence (BIAMA5_HUMAN) (Figure S9A, C). Previous studies have shown that the NTD of the Sin1 protein is important for its interaction with components of mTORC2, such as Rictor or MEKK2 (Chen et al., 2018; Cheng et al., 2005). The NTD-lacking isoform of Sin1, called Sin1δ the mTORC2 assembly and most of the mTORC2 functions (Yuan et al., 2015). Taken together, these results suggest that, although the precise function of NTD-lacking Sin1 is unclear, the variation of NTD-lacking Sin1 expression associated with the rs631287 genotype may underlie the risk of diverticular disease via mTORC2-independent mechanisms. Overall, these results emphasize that our puQTL analysis could provide new insights into associations between variants and complex traits, which were overlooked by previous eQTL analyses.

## Discussion

Here, we report a new computational framework for discovering genomic loci associated with promoter usage that was constructed by leveraging conventional RNA-seq datasets without additional costs. Our analysis provided insights into the regulation of promoter activity by short and long genomic variations. The combined analysis of transcript assembly and promoter activity estimates enabled us to identify the perturbation of “hidden” promoters, which were overlooked by most annotation-based analyses. In addition, our integrated analysis of puQTL with GWAS led to novel interpretations of the molecular mechanisms of complex traits. This study emphasizes the importance of puQTL analysis in understanding gene regulation and common disease pathogenesis. Our method requires only genotype and RNA-seq data and, thus, can be applied to other extensive datasets provided by international consortia, such as the GTEx project (The GTEx Consortium, 2020). Identifying puQTL in various tissues using the GTEx datasets will help elucidate the tissue-specific regulation of promoter activities by genomic variations and promoter-usage-associated disease mechanisms in clinically relevant tissues.

Changes in promoter usage can lead to changes in the relative abundance of transcripts harboring distinct 5′ untranslated regions (5′ UTR), which can affect mRNA translation rates (Leppek et al., 2018). In fact, the 5′ UTR includes various functional structures and elements, such as internal ribosome entry sites, binding sites for non-coding RNAs and RNA-binding proteins, RNA modification sites, hairpins, pseudo-knots, and RNA G-quadruplexes. These structures and elements regulate the recruitment of eukaryotic initiation factors to the 5′ end of mRNAs, and affect the mRNA translation rates, indicating the possibility that perturbation of promoter activity associated with genomic variations may result in changes in protein abundance. However, our analysis had a limitation: we focused exclusively on mRNA expression levels, rather than protein expression; therefore, we were unable to assess how promoter changes affect translation regulation. Previous studies have reported the association between genomic loci and protein expression levels (protein QTL: pQTL) (Battle et al., 2015; Sun et al., 2018; Wu et al., 2013); thus, by integrating the puQTL and pQTL data obtained from matched samples, we were able to discover promoter-change-associated protein expression changes. It is well known that protein expression levels are often uncorrelated with corresponding mRNA expression levels (Albert et al., 2014; Foss et al., 2007; Ghazalpour et al., 2011; Picotti et al., 2013), suggesting that it may become possible to interpret why protein levels change in the absence of a total mRNA expression change by focusing on promoter perturbation.

Promoter usage and splicing are closely interrelated. A recent study reported that splicing of internal exons promotes the activation of cryptic promoters, which is called exon-mediated activation of transcription starts (EMATS) (Fiszbein et al., 2019). Splicing factors interact with general TFs and recruit them to nearby weak promoters, resulting in promoter activation and an increase in transcriptional output (Fiszbein et al., 2019); however, the genetic factors that drive EMATS remain unexplored. In addition to gene expression, genomic variations also affect alternative splicing (splicing QTL: sQTL), thus implying that genomic variations affect the inclusion level of internal exons and lead to differential usage of nearby promoters in some genes. Although our analysis focused on the enrichment of genomic variations associated with promoter usage in promoters, enhancers, and TF binding sites (Figure 2), the combined analysis of puQTL and sQTL has the potential to identify genetic factors underlying splicing-mediated promoter activation. Our approach, which was developed to map puQTL using RNA-seq data, is valuable for exploring novel gene regulatory programs.

In this study, reference-guided transcript assembly using StringTie2 (Kovaka et al., 2019) and short-read RNA-seq reads of 438 samples enabled us to find “hidden” active promoters (Figure 3C, S7E, S8A), highlighting the fact that the standard gene annotations provided by Ensembl are incomplete. The precise construction of gene annotations is essential to explore the regulatory mechanism of promoter activity. However, the construction of transcript structures using only short-reads has been challenging. Notably, the StringTie2 software also works with long-read RNA sequencing reads and shows high performance in transcript assembly (Kovaka et al., 2019). A recent study generated a long-read RNA-seq dataset using the Oxford Nanopore Technologies platform from 88 samples from the GTEx project (Glinos et al., 2021). The leveraging of the GTEx short- and long-read RNA-seq dataset would improve our pipeline for mapping additional puQTL.

Taken together, the findings of this study suggest that our pipeline may be the current best practice for accurate RNA-seq-based puQTL mapping. Moving forward, by expanding this pipeline to extensive datasets in public databases, our strategy can help us further elucidate the genetic factors underlying transcriptional regulation and promote the novel discovery of the molecular mechanisms responsible for complex trait associations.

## Supplemental data

Supplemental Data include nine figures and one table.

## Acknowledgments

We thank all members of Mikita Suyama’s laboratory for their valuable discussions. The authors would like to thank Enago (www.enago.jp) for the English language review.

## Data and code availability

Genotype data of SNPs and indels of the 1000 Genomes Project is available at http://ftp.1000genomes.ebi.ac.uk/vol1/ftp/release/20130502/ and that of SVs is available at https://zenodo.org/record/5509980/files/SVs_1KGP_pgGTs.vcf.gz (Yan et al., 2021). RNA-seq data of the GEUVADIS Project is available at https://www.internationalgenome.org/data-portal/data-collection/geuvadis. The GTF file of Ensembl 104 transcript annotation is available at http://ftp.ensembl.org/pub/release-104/gtf/homo_sapiens/Homo_sapiens.GRCh38.104.gtf.gz. The TF footprints datasets for the GM12865 and GM06990 cell lines are available at https://resources.altius.org/~jvierstra/projects/footprinting.2020/per.dataset/, and we obtained “interval.all.fps.0.05.bed.gz” for each biosample.

The data of chromatin states for the GM12878 cell line is available at https://egg2.wustl.edu/roadmap/data/byFileType/chromhmmSegmentations/ChmmModels/imputed12marks/jointModel/final/E116_25_imputed12marks_hg38lift_mnemonics.bed.gz.

The recombination hotspots data is available at http://ftp.1000genomes.ebi.ac.uk/vol1/ftp/technical/working/20110106_recombination_hotspots/HapmapII_GRCh37_RecombinationHotspots.tar.gz.

A file of GWAS lead variants is available at https://www.ebi.ac.uk/gwas/docs/file-downloads, with the file name “gwas_catalog_v1.0.2-associations_e104_r2021-12-07.tsv.” Files of GWAS summary statistics are available at http://ftp.ebi.ac.uk/pub/databases/gwas/summary_statistics/GCST003001-GCST004000/GCST003043/harmonised/26192919-GCST003043-EFO_0003767.h.tsv.gz (IBD) and http://ftp.ebi.ac.uk/pub/databases/gwas/summary_statistics/GCST006001-GCST007000/GCST006479/clinical_c_K57.zip (diverticular disease). Results of QTL analyses are available on Zenodo (https://doi.org/10.5281/zenodo.6269326).

## References

1. Alasoo K, Rodrigues J, Danesh J, Freitag DF, Paul DS, Gaffney DJ. 2019. Genetic effects on promoter usage are highly context-specific and contribute to complex traits. Elife 8:e41673. doi:10.7554/elife.41673

2. Albert FW, Treusch S, Shockley AH, Bloom JS, Kruglyak L. 2014. Genetics of single-cell protein abundance variation in large yeast populations. Nature 506:494–497. doi:10.1038/nature12904

3. Battle A, Khan Z, Wang SH, Mitrano A, Ford MJ, Pritchard JK, Gilad Y. 2015. Impact of regulatory variation from RNA to protein. Science 347:664–667. doi:10.1126/science.1260793

4. Battle A, Mostafavi S, Zhu X, Potash JB, Weissman MM, McCormick C, Haudenschild CD, Beckman KB, Shi J, Mei R, Urban AE, Montgomery SB, Levinson DF, Koller D. 2013. Characterizing the genetic basis of transcriptome diversity through RNA-sequencing of 922 individuals. Genome Res 24:14–24. doi:10.1101/gr.155192.113

5. Bhattacharyya S, Chandra V, Vijayanand P, Ay F. 2019. Identification of significant chromatin contacts from HiChIP data by FitHiChIP. Nat Commun 10:4221. doi:10.1038/s41467-019-11950-y

6. Blauwendraat C, Francescatto M, Gibbs JR, Jansen IE, Simón-Sánchez J, Hernandez DG, Dillman AA, Singleton AB, Cookson MR, Rizzu P, Heutink P. 2016. Comprehensive promoter level expression quantitative trait loci analysis of the human frontal lobe. Genome Med 8:65. doi:10.1186/s13073-016-0320-1

7. Blum M, Chang H-Y, Chuguransky S, Grego T, Kandasaamy S, Mitchell A, Nuka G, Paysan-Lafosse T, Qureshi M, Raj S, Richardson L, Salazar GA, Williams L, Bork P, Bridge A, Gough J, Haft DH, Letunic I, Marchler-Bauer A, Mi H, Natale DA, Necci M, Orengo CA, Pandurangan AP, Rivoire C, Sigrist CJA, Sillitoe I, Thanki N, Thomas PD, Tosatto SCE, Wu CH, Bateman A, Finn RD. 2020. The InterPro protein families and domains database: 20 years on. Nucleic Acids Res 49:D344–D354. doi:10.1093/nar/gkaa977

8. Bray NL, Pimentel H, Melsted P, Pachter L. 2016. Near-optimal probabilistic RNA-seq quantification. Nat Biotechnol 34:525–527. doi:10.1038/nbt.3519

9. Chen L, Ge B, Casale FP, Vasquez L, Kwan T, Garrido-Martín D, Watt S, Yan Y, Kundu K, Ecker S, Datta A, Richardson D, Burden F, Mead D, Mann AL, Fernandez JM, Rowlston S, Wilder SP, Farrow S, Shao X, Lambourne JJ, Redensek A, Albers CA, Amstislavskiy V, Ashford S, Berentsen K, Bomba L, Bourque G, Bujold D, Busche S, Caron M, Chen S-H, Cheung W, Delaneau O, Dermitzakis ET, Elding H, Colgiu I, Bagger FO, Flicek P, Habibi E, Iotchkova V, Janssen-Megens E, Kim B, Lehrach H, Lowy E, Mandoli A, Matarese F, Maurano MT, Morris JA, Pancaldi V, Pourfarzad F, Rehnstrom K, Rendon A, Risch T, Sharifi N, Simon M-M, Sultan M, Valencia A, Walter K, Wang S-Y, Frontini M, Antonarakis SE, Clarke L, Yaspo M-L, Beck S, Guigo R, Rico D, Martens JHA, Ouwehand WH, Kuijpers TW, Paul DS, Stunnenberg HG, Stegle O, Downes K, Pastinen T, Soranzo N. 2016. Genetic Drivers of Epigenetic and Transcriptional Variation in Human Immune *Cell*s. Cell 167:1398–1414.e24. doi:10.1016/j.cell.2016.10.026

10. Chen S, Zhou Y, Chen Y, Gu J. 2018. fastp: an ultra-fast all-in-one FASTQ preprocessor. Bioinformatics 34:i884–i890. doi:10.1093/bioinformatics/bty560

11. Chen X, Liu M, Tian Y, Li J, Qi Y, Zhao D, Wu Z, Huang M, Wong CCL, Wang H-W, Wang J, Yang H, Xu Y. 2018. Cryo-EM structure of human mTOR complex 2. Cell Res 28:518–528. doi:10.1038/s41422-018-0029-3

12. Cheng J, Zhang D, Kim K, Zhao Yingxin, Zhao Yingming, Su B. 2005. Mip1, an MEKK2-Interacting Protein, Controls MEKK2 Dimerization and Activation. Mol Cell Biol 25:5955–5964. doi:10.1128/mcb.25.14.5955-5964.2005

13. Consortium IMSG, Consortium IIG, Liu JZ, van Sommeren S, Huang H, Ng SC, Alberts R, Takahashi A, Ripke S, Lee JC, Jostins L, Shah T, Abedian S, Cheon JH, Cho J, Daryani NE, Franke L, Fuyuno Y, Hart A, Juyal RC, Juyal G, Kim WH, Morris AP, Poustchi H, Newman WG, Midha V, Orchard TR, Vahedi H, Sood A, Sung JJY, Malekzadeh R, Westra H-J, Yamazaki K, Yang S-K, Barrett JC, Franke A, Alizadeh BZ, Parkes M, K TB, Daly MJ, Kubo M, Anderson CA, Weersma RK. 2015. Association analyses identify 38 susceptibility loci for inflammatory bowel disease and highlight shared genetic risk across populations. Nat Genet 47:979–986. doi:10.1038/ng.3359

14. Consortium RE, Kundaje A, Meuleman W, Ernst J, Bilenky M, Yen A, Heravi-Moussavi A, Kheradpour P, Zhang Z, Wang J, Ziller MJ, Amin V, Whitaker JW, Schultz MD, Ward LD, Sarkar A, Quon G, Sandstrom RS, Eaton ML, Wu Y-C, Pfenning AR, Wang X, Claussnitzer M, Liu Y, Coarfa C, Harris RA, Shoresh N, Epstein CB, Gjoneska E, Leung D, Xie W, Hawkins RD, Lister R, Hong C, Gascard P, Mungall AJ, Moore R, Chuah E, Tam A, Canfield TK, Hansen RS, Kaul R, Sabo PJ, Bansal MS, Carles A, Dixon JR, Farh K-H, Feizi S, Karlic R, Kim A-R, Kulkarni A, Li D, Lowdon R, Elliott G, Mercer TR, Neph SJ, Onuchic V, Polak P, Rajagopal N, Ray P, Sallari RC, Siebenthall KT, Sinnott-Armstrong NA, Stevens M, Thurman RE, Wu J, Zhang B, Zhou X, Beaudet AE, Boyer LA, Jager PLD, Farnham PJ, Fisher SJ, Haussler D, Jones SJM, Li W, Marra MA, McManus MT, Sunyaev S, Thomson JA, Tlsty TD, Tsai L-H, Wang W, Waterland RA, Zhang MQ, Chadwick LH, Bernstein BE, Costello JF, Ecker JR, Hirst M, Meissner A, Milosavljevic A, Ren B, Stamatoyannopoulos JA, Wang T, Kellis M. 2015. Integrative analysis of 111 reference human epigenomes. Nature 518:317. doi:10.1038/nature14248

15. Consortium T 1000 GP, Auton A, Brooks LD, Durbin RM, Garrison EP, Kang HM, Korbel JO, Marchini JL, McCarthy S, McVean GA, Abecasis GR. 2015. A global reference for human genetic variation. Nature 526:68. doi:10.1038/nature15393

16. Consortium TGte. 2020. The GTEx Consortium atlas of genetic regulatory effects across human tissues. Science 369:1318–1330. doi:10.1126/science.aaz1776

17. Delaneau O, Ongen H, Brown AA, Fort A, Panousis NI, Dermitzakis ET. 2017. A complete tool set for molecular QTL discovery and analysis. Nat Commun 8:15452. doi:10.1038/ncomms15452

18. Demircioğlu D, Cukuroglu E, Kindermans M, Nandi T, Calabrese C, Fonseca NA, Kahles A, Lehmann K-V, Stegle O, Brazma A, Brooks AN, Rätsch G, Tan P, Göke J. 2019. A Pan-cancer Transcriptome Analysis Reveals Pervasive Regulation through Alternative Promoters. Cell 178:1465–1477.e17. doi:10.1016/j.cell.2019.08.018

19. Dobin A, Davis CA, Schlesinger F, Drenkow J, Zaleski C, Jha S, Batut P, Chaisson M, Gingeras TR. 2013. STAR: ultrafast universal RNA-seq aligner. Bioinformatics 29:15–21. doi:10.1093/bioinformatics/bts635

20. Dunham I, Kundaje A, Aldred SF, Collins PJ, Davis CA, Doyle F, Epstein CB, Frietze S, Harrow J, Kaul R, Khatun J, Lajoie BR, Landt SG, Lee B-K, Pauli F, Rosenbloom KR, Sabo P, Safi A, Sanyal A, Shoresh N, Simon JM, Song L, Trinklein ND, Altshuler RC, Birney E, Brown JB, Cheng C, Djebali S, Dong X, Dunham I, Ernst J, Furey TS, Gerstein M, Giardine B, Greven M, Hardison RC, Harris RS, Herrero J, Hoffman MM, Iyer S, Kellis M, Khatun J, Kheradpour P, Kundaje A, Lassmann T, Li Q, Lin X, Marinov GK, Merkel A, Mortazavi A, Parker SCJ, Reddy TE, Rozowsky J, Schlesinger F, Thurman RE, Wang J, Ward LD, Whitfield TW, Wilder SP, Wu W, Xi HS, Yip KY, Zhuang J, Bernstein BE, Birney E, Dunham I, Green ED, Gunter C, Snyder M, Pazin MJ, Lowdon RF, Dillon LAL, Adams LB, Kelly CJ, Zhang J, Wexler JR, Green ED, Good PJ, Feingold EA, Bernstein BE, Birney E, Crawford GE, Dekker J, Elnitski L, Farnham PJ, Gerstein M, Giddings MC, Gingeras TR, Green ED, Guigó R, Hardison RC, Hubbard TJ, Kellis M, Kent WJ, Lieb JD, Margulies EH, Myers RM, Snyder M, Stamatoyannopoulos JA, Tenenbaum SA, Weng Z, White KP, Wold B, Khatun J, Yu Y, Wrobel J, Risk BA, Gunawardena HP, Kuiper HC, Maier CW, Xie L, Chen X, Giddings MC, Bernstein BE, Epstein CB, Shoresh N, Ernst J, Kheradpour P, Mikkelsen TS, Gillespie S, Goren A, Ram O, Zhang X, Wang L, Issner R, Coyne MJ, Durham T, Ku M, Truong T, Ward LD, Altshuler RC, Eaton ML, Kellis M, Djebali S, Davis CA, Merkel A, Dobin A, Lassmann T, Mortazavi A, Tanzer A, Lagarde J, Lin W, Schlesinger F, Xue C, Marinov GK, Khatun J, Williams BA, Zaleski C, Rozowsky J, Röder M, Kokocinski F, Abdelhamid RF, Alioto T, Antoshechkin I, Baer MT, Batut P, Bell I, Bell K, Chakrabortty S, Chen X, Chrast J, Curado J, Derrien T, Drenkow J, Dumais E, Dumais J, Duttagupta R, Fastuca M, Fejes-Toth K, Ferreira P, Foissac S, Fullwood MJ, Gao H, Gonzalez D, Gordon A, Gunawardena HP, Howald C, Jha S, Johnson R, Kapranov P, King B, Kingswood C, Li G, Luo OJ, Park E, Preall JB, Presaud K, Ribeca P, Risk BA, Robyr D, Ruan X, Sammeth M, Sandhu KS, Schaeffer L, See L-H, Shahab A, Skancke J, Suzuki AM, Takahashi H, Tilgner H, Trout D, Walters N, Wang Huaien, Wrobel J, Yu Y, Hayashizaki Y, Harrow J, Gerstein M, Hubbard TJ, Reymond A, Antonarakis SE, Hannon GJ, Giddings MC, Ruan Y, Wold B, Carninci P, Guigó R, Gingeras TR, Rosenbloom KR, Sloan CA, Learned K, Malladi VS, Wong MC, Barber GP, Cline MS, Dreszer TR, Heitner SG, Karolchik D, Kent WJ, Kirkup VM, Meyer LR, Long JC, Maddren M, Raney BJ, Furey TS, Song L, Grasfeder LL, Giresi PG, Lee B-K, Battenhouse A, Sheffield NC, Simon JM, Showers KA, Safi A, London D, Bhinge AA, Shestak C, Schaner MR, Kim SK, Zhang ZZ, Mieczkowski PA, Mieczkowska JO, Liu Z, McDaniell RM, Ni Y, Rashid NU, Kim MJ, Adar S, Zhang Zhancheng, Wang T, Winter D, Keefe D, Birney E, Iyer VR, Lieb JD, Crawford GE, Li G, Sandhu KS, Zheng M, Wang P, Luo OJ, Shahab A, Fullwood MJ, Ruan X, Ruan Y, Myers RM, Pauli F, Williams BA, Gertz J, Marinov GK, Reddy TE, Vielmetter J, Partridge E, Trout D, Varley KE, Gasper C, Bansal A, Pepke S, Jain P, Amrhein H, Bowling KM, Anaya M, Cross MK, King B, Muratet MA, Antoshechkin I, Newberry KM, McCue K, Nesmith AS, Fisher-Aylor KI, Pusey B, DeSalvo G, Parker SL, Balasubramanian Sreeram, Davis NS, Meadows SK, Eggleston T, Gunter C, Newberry JS, Levy SE, Absher DM, Mortazavi A, Wong WH, Wold B, Blow MJ, Visel A, Pennachio LA, Elnitski L, Margulies EH, Parker SCJ, Petrykowska HM, Abyzov A, Aken B, Barrell D, Barson G, Berry A, Bignell A, Boychenko V, Bussotti G, Chrast J, Davidson C, Derrien T, Despacio-Reyes G, Diekhans M, Ezkurdia I, Frankish A, Gilbert J, Gonzalez JM, Griffiths E, Harte R, Hendrix DA, Howald C, Hunt T, Jungreis I, Kay M, Khurana E, Kokocinski F, Leng J, Lin MF, Loveland J, Lu Z, Manthravadi D, Mariotti M, Mudge J, Mukherjee G, Notredame C, Pei B, Rodriguez JM, Saunders G, Sboner A, Searle S, Sisu C, Snow C, Steward C, Tanzer A, Tapanari E, Tress ML, van Baren MJ, Walters N, Washietl S, Wilming L, Zadissa A, Zhang Zhengdong, Brent M, Haussler D, Kellis M, Valencia A, Gerstein M, Reymond A, Guigó R, Harrow J, Hubbard TJ, Landt SG, Frietze S, Abyzov A, Addleman N, Alexander RP, Auerbach RK, Balasubramanian Suganthi, Bettinger K, Bhardwaj N, Boyle AP, Cao AR, Cayting P, Charos A, Cheng Y, Cheng C, Eastman C, Euskirchen G, Fleming JD, Grubert F, Habegger L, Hariharan M, Harmanci A, Iyengar S, Jin VX, Karczewski KJ, Kasowski M, Lacroute P, Lam H, Lamarre-Vincent N, Leng J, Lian J, Lindahl-Allen M, Min R, Miotto B, Monahan H, Moqtaderi Z, Mu XJ, O’Geen H, Ouyang Z, Patacsil D, Pei B, Raha D, Ramirez L, Reed B, Rozowsky J, Sboner A, Shi M, Sisu C, Slifer T, Witt H, Wu L, Xu X, Yan K-K, Yang X, Yip KY, Zhang Zhengdong, Struhl K, Weissman SM, Gerstein M, Farnham PJ, Snyder M, Tenenbaum SA, Penalva LO, Doyle F, Karmakar S, Landt SG, Bhanvadia RR, Choudhury A, Domanus M, Ma L, Moran J, Patacsil D, Slifer T, Victorsen A, Yang X, Snyder M, White KP, Auer T, Centanin L, Eichenlaub M, Gruhl F, Heermann S, Hoeckendorf B, Inoue D, Kellner T, Kirchmaier S, Mueller C, Reinhardt R, Schertel L, Schneider S, Sinn R, Wittbrodt B, Wittbrodt J, Weng Z, Whitfield TW, Wang J, Collins PJ, Aldred SF, Trinklein ND, Partridge EC, Myers RM, Dekker J, Jain G, Lajoie BR, Sanyal A, Balasundaram G, Bates DL, Byron R, Canfield TK, Diegel MJ, Dunn D, Ebersol AK, Frum T, Garg K, Gist E, Hansen RS, Boatman L, Haugen E, Humbert R, Jain G, Johnson AK, Johnson EM, Kutyavin TV, Lajoie BR, Lee K, Lotakis D, Maurano MT, Neph SJ, Neri FV, Nguyen ED, Qu H, Reynolds AP, Roach V, Rynes E, Sabo P, Sanchez ME, Sandstrom RS, Sanyal A, Shafer AO, Stergachis AB, Thomas S, Thurman RE, Vernot B, Vierstra J, Vong S, Wang Hao, Weaver MA, Yan Y, Zhang M, Akey JM, Bender M, Dorschner MO, Groudine M, MacCoss MJ, Navas P, Stamatoyannopoulos G, Kaul R, Dekker J, Stamatoyannopoulos JA, Dunham I, Beal K, Brazma A, Flicek P, Herrero J, Johnson N, Keefe D, Lukk M, Luscombe NM, Sobral D, Vaquerizas JM, Wilder SP, Batzoglou S, Sidow A, Hussami N, Kyriazopoulou-Panagiotopoulou S, Libbrecht MW, Schaub MA, Kundaje A, Hardison RC, Miller W, Giardine B, Harris RS, Wu W, Bickel PJ, Banfai B, Boley NP, Brown JB, Huang H, Li Q, Li JJ, Noble WS, Bilmes JA, Buske OJ, Hoffman MM, Sahu AD, Kharchenko PV, Park PJ, Baker D, Taylor J, Weng Z, Iyer S, Dong X, Greven M, Lin X, Wang J, Xi HS, Zhuang J, Gerstein M, Alexander RP, Balasubramanian Suganthi, Cheng C, Harmanci A, Lochovsky L, Min R, Mu XJ, Rozowsky J, Yan K-K, Yip KY, Birney E. 2012. An integrated encyclopedia of DNA elements in the human genome. Nature 489:nature11247. doi:10.1038/nature11247

21. Edgar RC. 2004. MUSCLE: multiple sequence alignment with high accuracy and high throughput. Nucleic Acids Res 32:1792–1797. doi:10.1093/nar/gkh340

22. Ernst J, Kellis M. 2017. Chromatin-state discovery and genome annotation with ChromHMM. Nat Protoc 12:2478. doi:10.1038/nprot.2017.124

23. Fiszbein A, Krick KS, Begg BE, Burge CB. 2019. Exon-Mediated Activation of Transcription Starts. Cell. doi:10.1016/j.cell.2019.11.002

24. Foss EJ, Radulovic D, Shaffer SA, Ruderfer DM, Bedalov A, Goodlett DR, Kruglyak L. 2007. Genetic basis of proteome variation in yeast. Nat Genet 39:1369–1375. doi:10.1038/ng.2007.22

25. Garieri M, Delaneau O, Santoni F, Fish RJ, Mull D, Carninci P, Dermitzakis ET, Antonarakis SE, Fort A. 2017. The effect of genetic variation on promoter usage and enhancer activity. Nat Commun 8:1358. doi:10.1038/s41467-017-01467-7

26. Ghazalpour A, Bennett B, Petyuk VA, Orozco L, Hagopian R, Mungrue IN, Farber CR, Sinsheimer J, Kang HM, Furlotte N, Park CC, Wen P-Z, Brewer H, Weitz K, Camp DG, Pan C, Yordanova R, Neuhaus I, Tilford C, Siemers N, Gargalovic P, Eskin E, Kirchgessner T, Smith DJ, Smith RD, Lusis AJ. 2011. Comparative Analysis of Proteome and Transcriptome Variation in Mouse. Plos Genet 7:e1001393. doi:10.1371/journal.pgen.1001393

27. Glinos DA, Garborcauskas G, Hoffman P, Ehsan N, Jiang L, Gokden A, Dai X, Aguet F, Brown KL, Garimella K, Bowers T, Costello M, Ardlie K, Jian R, Tucker NR, Ellinor PT, Harrington ED, Tang H, Snyder M, Juul S, Mohammadi P, MacArthur DG, Lappalainen T, Cummings B. 2021. Transcriptome variation in human tissues revealed by long-read sequencing. bioRxiv. doi:10.1101/2021.01.22.427687

28. Haeussler M, Zweig AS, Tyner C, Speir ML, Rosenbloom KR, Raney BJ, Lee CM, Lee BT, Hinrichs AS, Gonzalez JN, Gibson D, Diekhans M, Clawson H, Casper J, Barber GP, Haussler D, Kuhn RM, Kent WJ. 2019. The UCSC Genome Browser database: 2019 update. Nucleic Acids Res 47:D853–D858. doi:10.1093/nar/gky1095

29. (IIBDGC) TIIGC, Jostins L, Ripke S, Weersma RK, Duerr RH, McGovern DP, Hui KY, Lee JC, Schumm LP, Sharma Y, Anderson CA, Essers J, Mitrovic M, Ning K, Cleynen I, Theatre E, Spain SL, Raychaudhuri S, Goyette P, Wei Z, Abraham C, Achkar J-P, Ahmad T, Amininejad L, Ananthakrishnan AN, Andersen V, Andrews JM, Baidoo L, Balschun T, Bampton PA, Bitton A, Boucher G, Brand S, Büning C, Cohain A, Cichon S, D’Amato M, Jong DD, Devaney KL, Dubinsky M, Edwards C, Ellinghaus D, Ferguson LR, Franchimont D, Fransen K, Gearry R, Georges M, Gieger C, Glas J, Haritunians T, Hart A, Hawkey C, Hedl M, Hu X, Karlsen TH, Kupcinskas L, Kugathasan S, Latiano A, Laukens D, Lawrance IC, Lees CW, Louis E, Mahy G, Mansfield J, Morgan AR, Mowat C, Newman W, Palmieri O, Ponsioen CY, Potocnik U, Prescott NJ, Regueiro M, Rotter JI, Russell RK, Sanderson JD, Sans M, Satsangi J, Schreiber S, Simms LA, Sventoraityte J, Targan SR, Taylor KD, Tremelling M, Verspaget HW, Vos MD, Wijmenga C, Wilson DC, Winkelmann J, Xavier RJ, Zeissig S, Zhang B, Zhang CK, Zhao H, Silverberg MS, Annese V, Hakonarson H, Brant SR, Radford-Smith G, Mathew CG, Rioux JD, Schadt EE, Daly MJ, Franke A, Parkes M, Vermeire S, Barrett JC, Cho JH. 2012. Host-microbe interactions have shaped the genetic architecture of inflammatory bowel disease. Nature 491:119–124. doi:10.1038/nature11582

30. Jacinto E, Facchinetti V, Liu D, Soto N, Wei S, Jung SY, Huang Q, Qin J, Su B. 2006. SIN1/MIP1 Maintains rictor-mTOR Complex Integrity and Regulates Akt Phosphorylation and Substrate Specificity. Cell 127:125–137. doi:10.1016/j.cell.2006.08.033

31. Kovaka S, Zimin AV, Pertea GM, Razaghi R, Salzberg SL, Pertea M. 2019. Transcriptome assembly from long-read RNA-seq alignments with StringTie2. Genome Biol 20:278. doi:10.1186/s13059-019-1910-1

32. de Lange KM, Moutsianas L, Lee JC, Lamb CA, Luo Y, Kennedy NA, Jostins L, Rice DL, Gutierrez-Achury J, Ji S-G, Heap G, Nimmo ER, Edwards C, Henderson P, Mowat C, Sanderson J, Satsangi J, Simmons A, Wilson DC, Tremelling M, Hart A, Mathew CG, Newman WG, Parkes M, Lees CW, Uhlig H, Hawkey C, Prescott NJ, Ahmad T, Mansfield JC, Anderson CA, Barrett JC. 2017. Genome-wide association study implicates immune activation of multiple integrin genes in inflammatory bowel disease. Nat Genet 49: 256–261. doi:10.1038/ng.3760

33. Lappalainen T, Sammeth M, Friedländer MR, Hoen PAC’t, Monlong J, Rivas MA, Gonzàlez-Porta M, Kurbatova N, Griebel T, Ferreira PG, Barann M, Wieland T, Greger L, van Iterson M, Almlöf J, Ribeca P, Pulyakhina I, Esser D, Giger T, Tikhonov A, Sultan M, Bertier G, MacArthur DG, Lek M, Lizano E, Buermans HPJ, Padioleau I, Schwarzmayr T, Karlberg O, Ongen H, Kilpinen H, Beltran S, Gut M, Kahlem K, Amstislavskiy V, Stegle O, Pirinen M, Montgomery SB, Donnelly P, McCarthy MI, Flicek P, Strom TM, Consortium G, Lehrach H, Schreiber S, Sudbrak R, Carracedo A, Antonarakis SE, Häsler R, Syvänen A-C, van Ommen G-J, Brazma A, Meitinger T, Rosenstiel P, Guigó R, Gut IG, Estivill X, Dermitzakis ET. 2013. Transcriptome and genome sequencing uncovers functional variation in humans. Nature 501:506–11. doi:10.1038/nature12531

34. Leppek K, Das R, Barna M. 2018. Functional 5′ UTR mRNA structures in eukaryotic translation regulation and how to find them. Nat Rev Mol Cell Bio 19:158–174. doi:10.1038/nrm.2017.103

35. Liao Y, Smyth GK, Shi W. 2014. featureCounts: an efficient general purpose program for assigning sequence reads to genomic features. Bioinformatics 30:923–930. doi:10.1093/bioinformatics/btt656

36. Lisak DA, Schacht T, Enders V, Habicht J, Kiviluoto S, Schneider J, Henke N, Bultynck G, Methner A. 2015. The transmembrane Bax inhibitor motif (TMBIM) containing protein family: Tissue expression, intracellular localization and effects on the ER CA2+-filling state. Biochimica Et Biophysica Acta Bba - Mol Cell Res 1853:2104–2114. doi:10.1016/j.bbamcr.2015.03.002

37. Luzi L, Confalonieri S, Fiore PPD, Pelicci PG. 2000. Evolution of Shc functions from nematode to human. Curr Opin Genet Dev 10:668–674. doi:10.1016/s0959-437x(00)00146-5

38. MacArthur J, Bowler E, Cerezo M, Gil L, Hall P, Hastings E, Junkins H, McMahon A, Milano A, Morales J, Pendlington ZM, Welter D, Burdett T, Hindorff L, Flicek P, Cunningham F, Parkinson H. 2017. The new NHGRI-EBI Catalog of published genome-wide association studies (GWAS Catalog). Nucleic Acids Res 45:D896–D901. doi:10.1093/nar/gkw1133

39. Maguire LH, Handelman SK, Du X, Chen Y, Pers TH, Speliotes EK. 2018. Genome-wide association analyses identify 39 new susceptibility loci for diverticular disease. Nat Genet 50:1359–1365. doi:10.1038/s41588-018-0203-z

40. Mirdita M, Schütze K, Moriwaki Y, Heo L, Ovchinnikov S, Steinegger M. 2021. ColabFold - Making protein folding accessible to all. Biorxiv 2021.08.15.456425. doi:10.1101/2021.08.15.456425

41. Mumbach MR, Satpathy AT, Boyle EA, Dai C, Gowen BG, Cho SW, Nguyen ML, Rubin AJ, Granja JM, Kazane KR, Wei Y, Nguyen T, Greenside PG, Corces MR, Tycko J, Simeonov DR, Suliman N, Li R, Xu J, Flynn RA, Kundaje A, Khavari PA, Marson A, Corn JE, Quertermous T, Greenleaf WJ, Chang HY. 2017. Enhancer connectome in primary human cells identifies target genes of disease-associated DNA elements. Nat Genet 49:1602–1612. doi:10.1038/ng.3963

42. Nica AC, Dermitzakis ET. 2013. Expression quantitative trait loci: present and future. Philosophical Transactions Royal Soc B Biological Sci 368:20120362. doi:10.1098/rstb.2012.0362

43. Nica AC, Montgomery SB, Dimas AS, Stranger BE, Beazley C, Barroso I, Dermitzakis ET. 2010. Candidate Causal Regulatory Effects by Integration of Expression QTLs with Complex Trait Genetic Associations. Plos Genet 6:e1000895. doi:10.1371/journal.pgen.1000895

44. Nozell S, Chen X. 2002. p21B, a variant of p21Waf1/Cip1, is induced by the p53 family. Oncogene 21:1285–1294. doi:10.1038/sj.onc.1205191

45. Ouelle DE, Zindy F, Ashmun RA, Sherr CJ. 1995. Alternative reading frames of the INK4a tumor suppressor gene encode two unrelated proteins capable of inducing cell cycle arrest. Cell 83:993– 1000. doi:10.1016/0092-8674(95)90214-7

46. Patro R, Duggal G, Love MI, Irizarry RA, Kingsford C. 2017. Salmon provides fast and bias-aware quantification of transcript expression. Nat Methods 14:417–419. doi:10.1038/nmeth.4197

47. Picotti P, Clément-Ziza M, Lam H, Campbell DS, Schmidt A, Deutsch EW, Röst H, Sun Z, Rinner O, Reiter L, Shen Q, Michaelson JJ, Frei A, Alberti S, Kusebauch U, Wollscheid B, Moritz RL, Beyer A, Aebersold R. 2013. A complete mass-spectrometric map of the yeast proteome applied to quantitative trait analysis. Nature 494:266–270. doi:10.1038/nature11835

48. Pozniak CD, Radinovic S, Yang A, McKeon F, Kaplan DR, Miller FD. 2000. An Anti-Apoptotic Role for the p53 Family Member, p73, During Developmental Neuron Death. Science 289:304–306. doi:10.1126/science.289.5477.304

49. Purcell S, Neale B, Todd-Brown K, Thomas L, Ferreira MAR, Bender D, Maller J, Sklar P, de Bakker PIW, Daly MJ, Sham PC. 2007. PLINK: A Tool Set for Whole-Genome Association and Population-Based Linkage Analyses. Am J Hum Genetics 81:559–575. doi:10.1086/519795

50. Ramírez F, Ryan DP, Grüning B, Bhardwaj V, Kilpert F, Richter AS, Heyne S, Dündar F, Manke T. 2016. deepTools2: a next generation web server for deep-sequencing data analysis. Nucleic Acids Res 44:W160–W165. doi:10.1093/nar/gkw257

51. Richards AL, Watza D, Findley A, Alazizi A, Wen X, Pai AA, Pique-Regi R, Luca F. 2017. Environmental perturbations lead to extensive directional shifts in RNA processing. Plos Genet 13:e1006995. doi:10.1371/journal.pgen.1006995

52. Robinson MD, McCarthy DJ, Smyth GK. 2010. edgeR: a Bioconductor package for differential expression analysis of digital gene expression data. Bioinformatics 26:139–140. doi:10.1093/bioinformatics/btp616

53. Schor IE, Degner JF, Harnett D, Cannavò E, Casale FP, Shim H, Garfield DA, Birney E, Stephens M, Stegle O, Furlong EEM. 2017. Promoter shape varies across populations and affects promoter evolution and expression noise. Nat Genet 49:550–558. doi:10.1038/ng.3791

54. Servant N, Varoquaux N, Lajoie BR, Viara E, Chen C-J, Vert J-P, Heard E, Dekker J, Barillot E. 2015. HiC-Pro: an optimized and flexible pipeline for Hi-C data processing. Genome Biol 16:259. doi:10.1186/s13059-015-0831-x

55. Shen W, Le S, Li Y, Hu F. 2016. SeqKit: A Cross-Platform and Ultrafast Toolkit for FASTA/Q File Manipulation. Plos One 11:e0163962. doi:10.1371/journal.pone.0163962

56. Sherr CJ. 2001. The INK4a/ARF network in tumour suppression. Nat Rev Mol Cell Bio 2:731–737. doi:10.1038/35096061

57. Shukla S, Fujita K, Xiao Q, Liao Z, Garfield S, Srinivasula SM. 2011. A shear stress responsive gene product PP1201 protects against Fas-mediated apoptosis by reducing Fas expression on the cell surface. Apoptosis 16:162–173. doi:10.1007/s10495-010-0556-y

58. Srivastava A, Malik L, Sarkar H, Zakeri M, Almodaresi F, Soneson C, Love MI, Kingsford C, Patro R. 2020. Alignment and mapping methodology influence transcript abundance estimation. Genome Biol 21:239. doi:10.1186/s13059-020-02151-8

59. Stegle O, Parts L, Piipari M, Winn J, Durbin R. 2012. Using probabilistic estimation of expression residuals (PEER) to obtain increased power and interpretability of gene expression analyses. Nat Protoc 7:500–507. doi:10.1038/nprot.2011.457

60. Sun BB, Maranville JC, Peters JE, Stacey D, Staley JR, Blackshaw J, Burgess S, Jiang T, Paige E, Surendran P, Oliver-Williams C, Kamat MA, Prins BP, Wilcox SK, Zimmerman ES, Chi A, Bansal N, Spain SL, Wood AM, Morrell NW, Bradley JR, Janjic N, Roberts DJ, Ouwehand WH, Todd JA, Soranzo N, Suhre K, Paul DS, Fox CS, Plenge RM, Danesh J, Runz H, Butterworth AS. 2018. Genomic atlas of the human plasma proteome. Nature 558:73–79. doi:10.1038/s41586-018-0175-2

61. Ventura A, Luzi L, Pacini S, Baldari CT, Pelicci PG. 2002. The p66Shc Longevity Gene Is Silenced through Epigenetic Modifications of an Alternative Promoter*. J Biol Chem 277:22370–22376. doi:10.1074/jbc.m200280200

62. Vierstra J, Lazar J, Sandstrom R, Halow J, Lee K, Bates D, Diegel M, Dunn D, Neri F, Haugen E, Rynes E, Reynolds A, Nelson J, Johnson A, Frerker M, Buckley M, Kaul R, Meuleman W, Stamatoyannopoulos JA. 2020. Global reference mapping of human transcription factor footprints. Nature 583:729–736. doi:10.1038/s41586-020-2528-x

63. Wallace C. 2021. A more accurate method for colocalisation analysis allowing for multiple causal variants. Plos Genet 17:e1009440. doi:10.1371/journal.pgen.1009440

64. Wang G, Sarkar A, Carbonetto P, Stephens M. 2020. A simple new approach to variable selection in regression, with application to genetic fine mapping. J Royal Statistical Soc Ser B Statistical Methodol 82:1273–1300. doi:10.1111/rssb.12388

65. Wu DC, Yao J, Ho KS, Lambowitz AM, Wilke CO. 2018. Limitations of alignment-free tools in total RNA-seq quantification. Bmc Genomics 19:510. doi:10.1186/s12864-018-4869-5

66. Wu L, Candille SI, Choi Y, Xie D, Jiang L, Li-Pook-Than J, Tang H, Snyder M. 2013. Variation and genetic control of protein abundance in humans. Nature 499:79–82. doi:10.1038/nature12223

67. Yan SM, Sherman RM, Taylor DJ, Nair DR, Bortvin AN, Schatz MC, McCoy RC. 2021. Local adaptation and archaic introgression shape global diversity at human structural variant loci. Elife 10:e67615. doi:10.7554/elife.67615

68. Yang A, Kaghad M, Wang Y, Gillett E, Fleming MD, Dötsch V, Andrews NC, Caput D, McKeon F. 1998. p63, a p53 Homolog at 3q27–29, Encodes Multiple Products with Transactivating, Death-Inducing, and Dominant-Negative Activities. Mol Cell 2:305–316. doi:10.1016/s1097-2765(00)80275-0

69. Yang Q, Inoki K, Ikenoue T, Guan K-L. 2006. Identification of Sin1 as an essential TORC2 component required for complex formation and kinase activity. Gene Dev 20:2820–2832. doi:10.1101/gad.1461206

70. Yuan Y, Pan B, Sun H, Chen G, Su B, Huang Y. 2015. Characterization of Sin1 Isoforms Reveals an mTOR-Dependent and Independent Function of Sin1γ doi:10.1371/journal.pone.0135017

71. Zaika AI, Slade N, Erster SH, Sansome C, Joseph TW, Pearl M, Chalas E, Moll UM. 2002. ΔDominant-Negative Inhibitor of Wild-type p53 and TAp73, Is Up-regulated in Human Tumors. J Exp Medicine 196:765–780. doi:10.1084/jem.20020179

72. Zhang Y, Liu T, Meyer CA, Eeckhoute J, Johnson DS, Bernstein BE, Nusbaum C, Myers RM, Brown M, Li W, Liu XS. 2008. Model-based Analysis of ChIP-Seq (MACS). Genome Biol 9:R137. doi:10.1186/gb-2008-9-9-r137

73. Zhao H, Ito A, Sakai N, Matsuzawa Y, Yamashita S, Nojima H. 2006. RECS1 is a Negative Regulator of Matrix Metalloproteinase-9 Production and Aged RECS1 Knockout Mice are Prone to Aortic Dilation. Circ J 70:615–624. doi:10.1253/circj.70.615

74. Zou Y, Carbonetto P, Wang G, Stephens M. 2021. Fine-mapping from summary data with the “Sum of Single Effects” model. Biorxiv 2021.11.03.467167. doi:10.1101/2021.11.03.467167

