## Supplemental Figure 1-9 for "Mapping of promoter usage QTL using RNA-seq data reveals their contributions to complex traits"

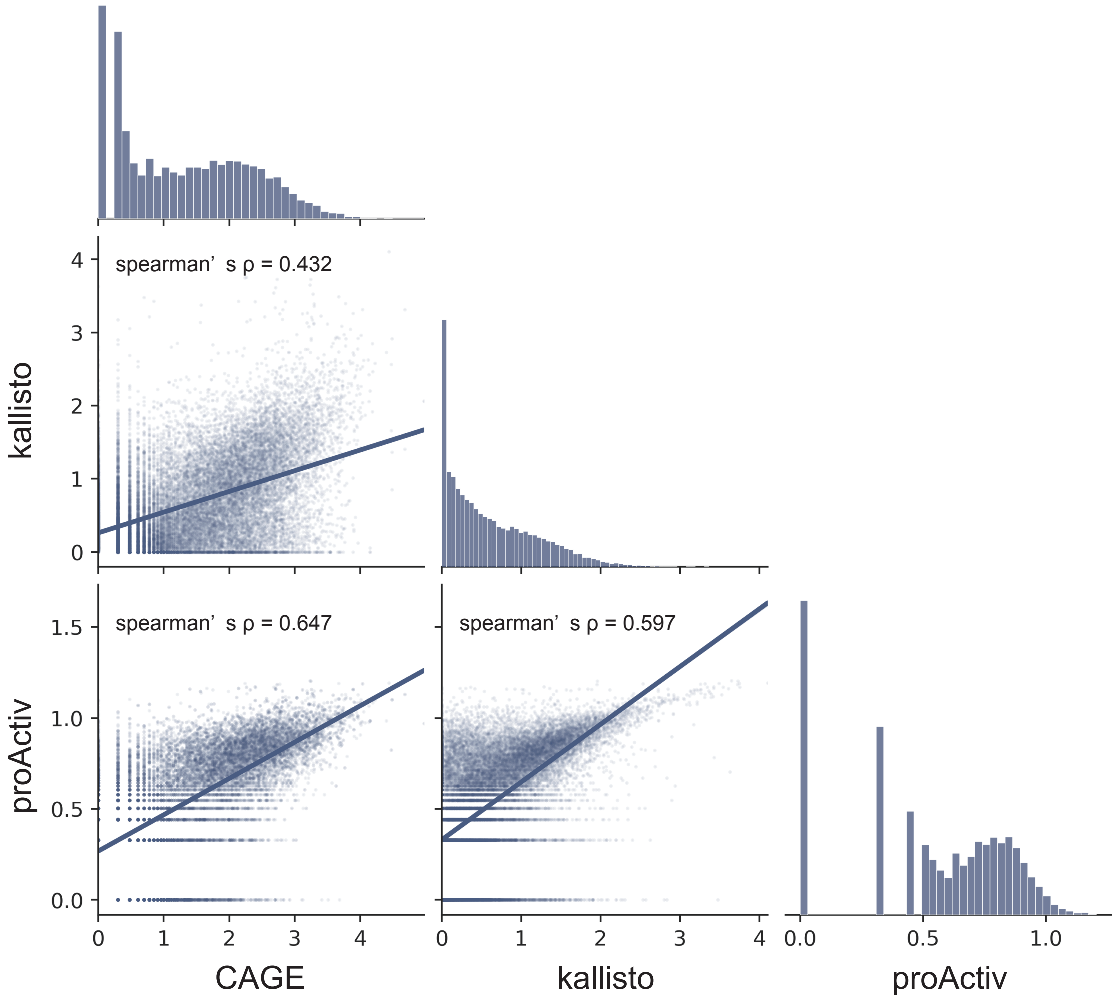


**Supplemental Figure 1. Correlation of estimated promoter activites among three different methods.** Activity scores of each promoter of the GM12878 cell line are plotted with regression lines.

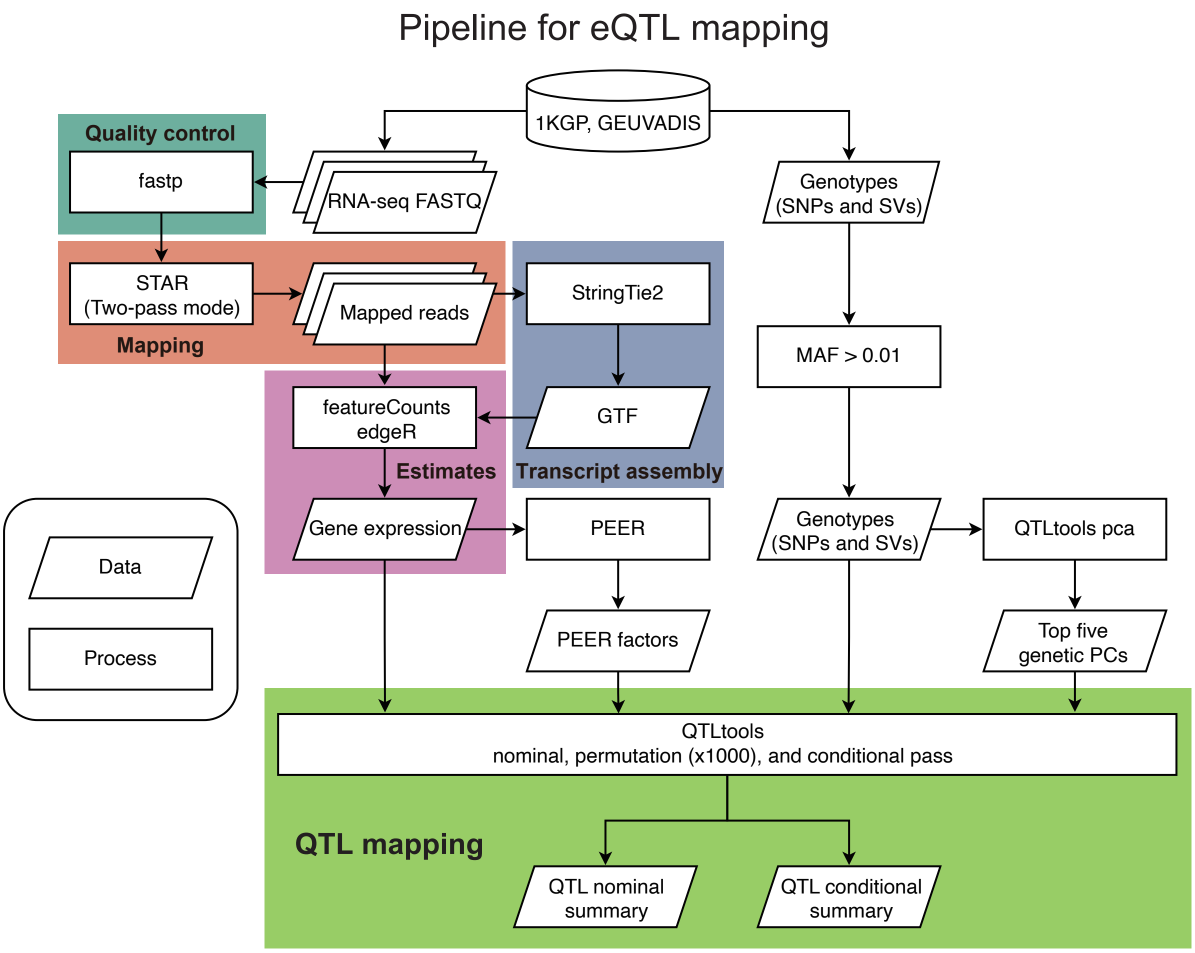


**Supplemental Figure 2. Pipeline for eQTL mapping.** 1KGP, the 1000 Genomes Project; MAF, Minor Allele Frequency; GTF, Gene Transfer Format.

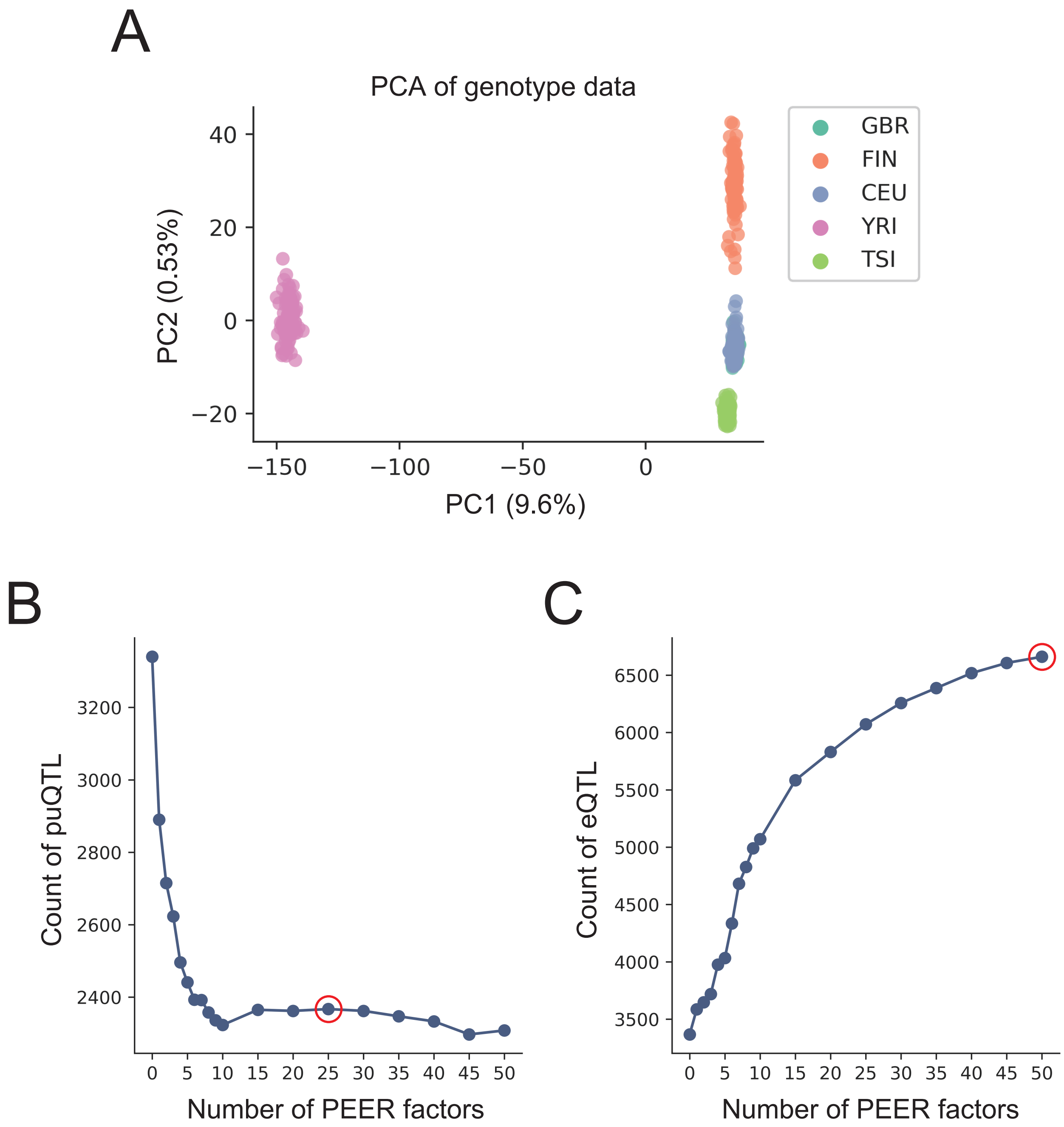


**Supplemental Figure 3. Covariates for QTL analysis.** (A) Projection on the first two principal components of the normalized genotype data, labelled for populations. GBR, British in England and Scotland; FIN, Finnish in Finland; CEU, Utah residents (CEPH) with Northern and Western European ancestry; YRI, Yoruba in Ibadan, Nigeria; TSI, Toscani in Italia. (B, C) The number of puQTL (B) and eQTL (C) (nominal $P < 1.0 \times 10^{-5}$) identified (y-axis) versus the number of PEER factors used (x-axis). Red circles represent the number of PEER factors used for the following analysis.

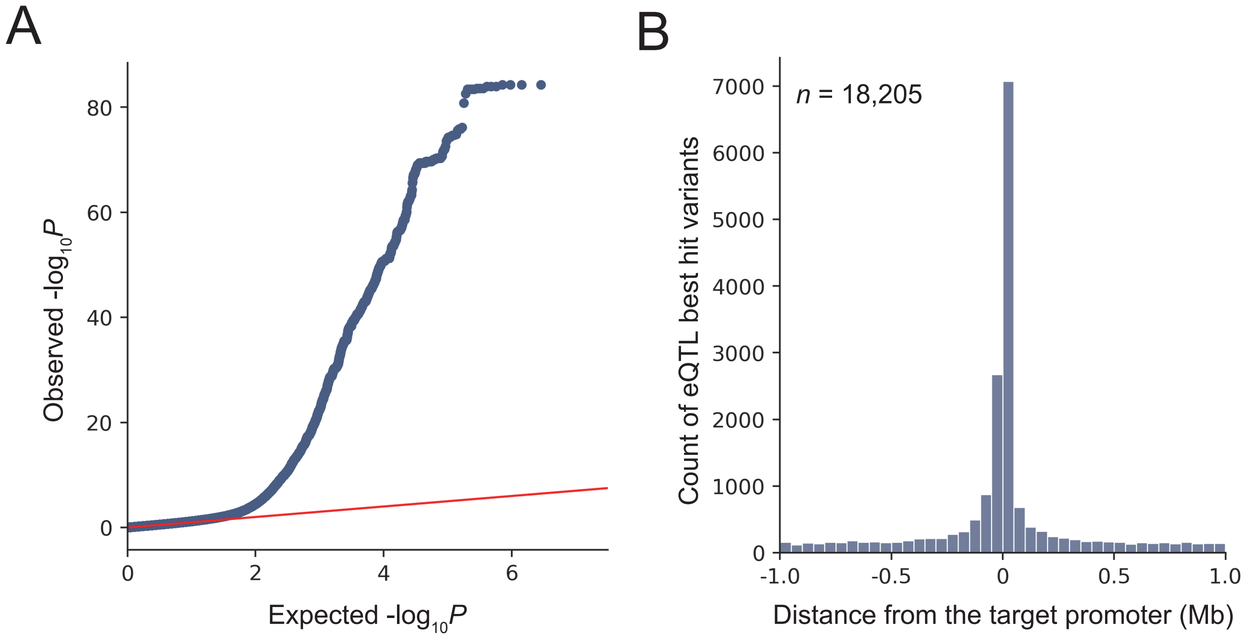


**Supplemental Figure 4. eQTL analysis results.** (A) Quantile–quantile plot of *P*-values. The nominal pass results of chromosome 22 are plotted and a red line indicates expected *P*-values under the null hypothesis. (B) Distribution of the distance of eQTL best hit variants from the target promoters.

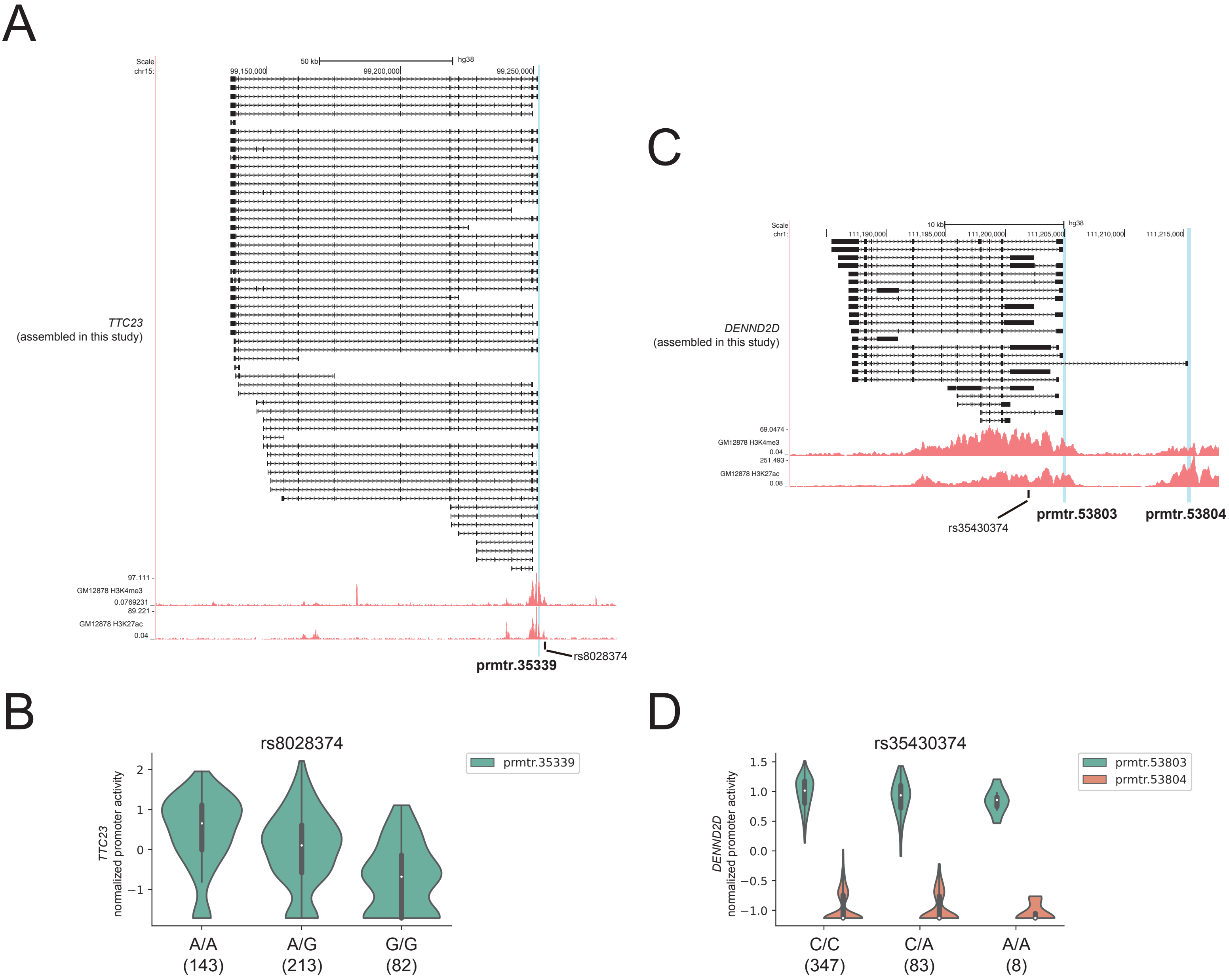


**Supplemental Figure 5. Examples of puQTL.** (A) The *TTC23* gene locus. Structures of the *TTC23* assembled in this study are shown with ENCODE GM12878 H3K4me3 and H3K27ac ChIP-seq signals. A vertical blue bar indicate the location of an active promoter, prmtr.35339. A black bar indicates the location of a variant rs8028374. (B) Comparison of the promoter activities of the *TTC23* gene among rs8028374 genotypes. The numbers in parentheses indicate sample size. (C) The *DENND2D* gene locus. (D) Comparison of the promoter activities of the *DENND2D* gene among rs35430374 genotypes.

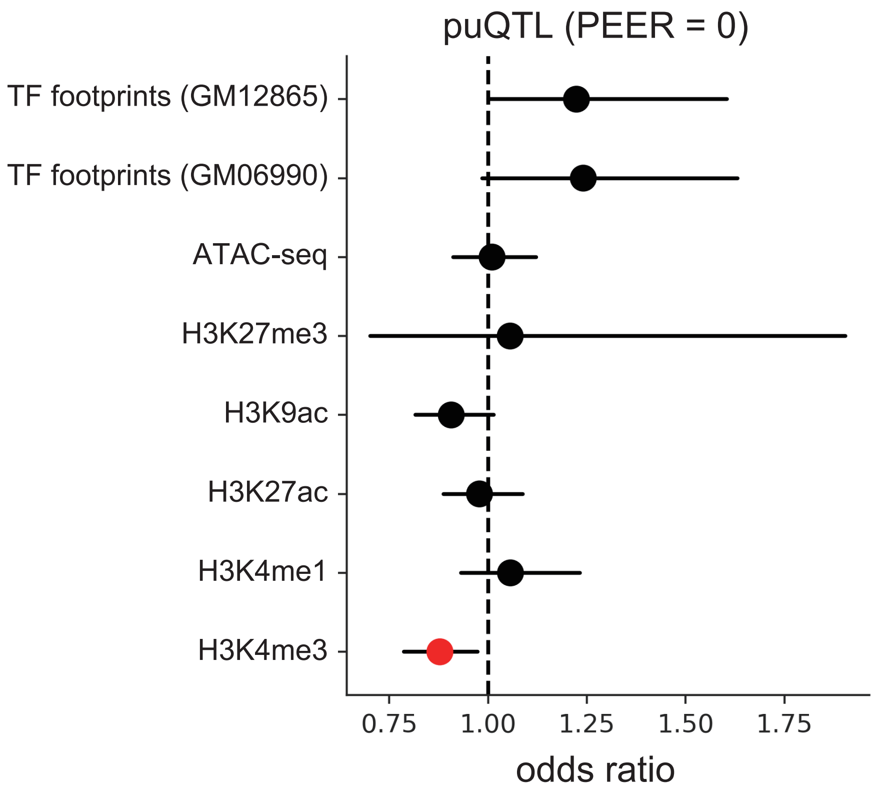


**Supplemental Figure 6. Enrichment of puQTL identified without PEER factors in epigenetic features.** Enrichment of peaks of histone mark ChIP-seq and transcription factor footprints. Red dots represent significant enrichment at the 5% FDR level and bars show 95% confidence intervals.

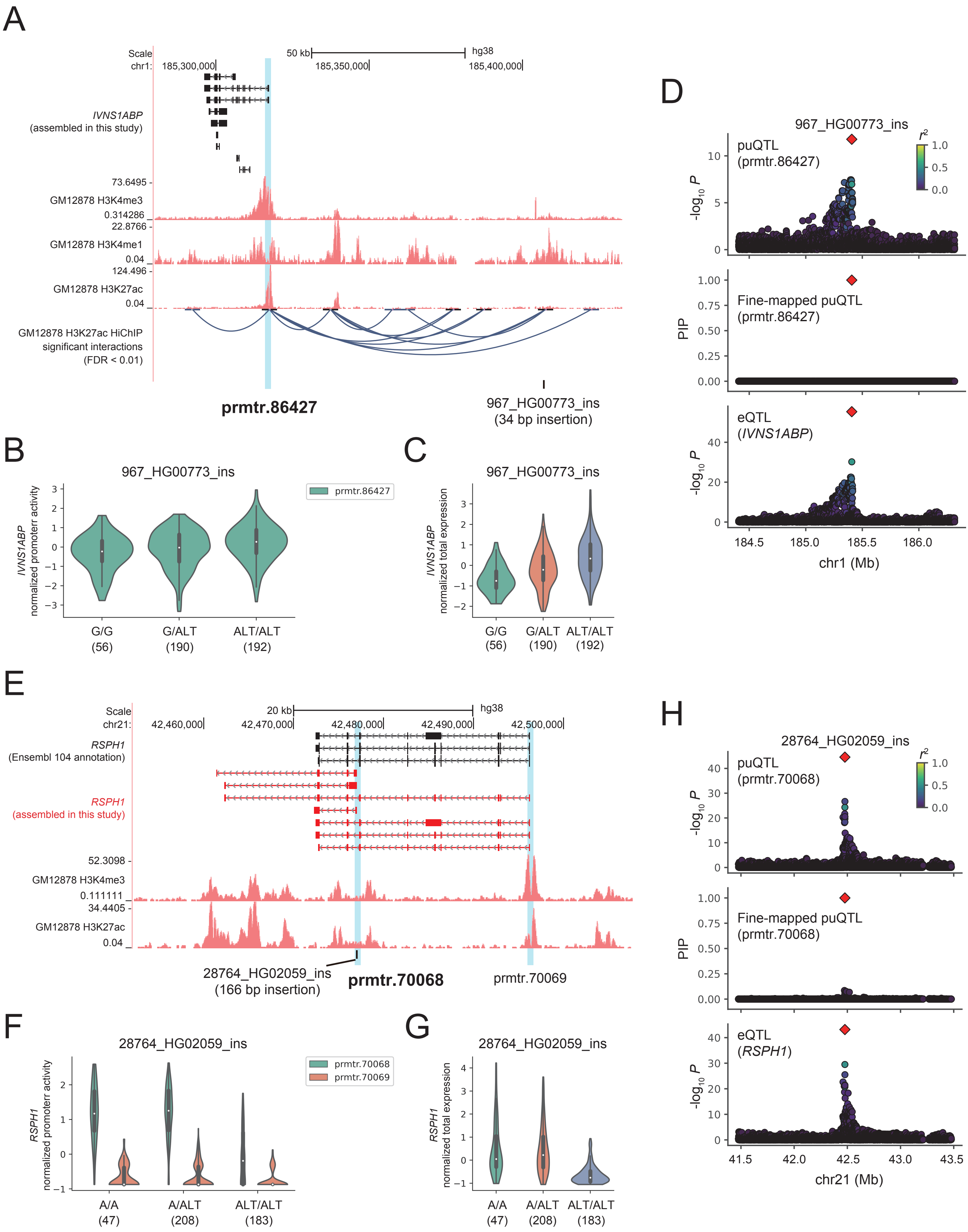
A
Scale
chr1:
50 kb
hg38
185,300,000
185,350,000
185,400,000
IVNS1ABP
(assembled in this study)
73.6495
GM12878 H3K4me3
0.314286
22.8766
GM12878 H3K4me1
0.04
124.496
GM12878 H3K27ac
0.04
GM12878 H3K27ac HiChIP
significant interactions
(FDR < 0.01)
prmtr.86427
967_HG00773_ins
(34 bp insertion)
B
967_HG00773_ins
IVNS1ABP normalized promoterr activity
prmtr.86427
G/G (56)
G/ALT (190)
ALT/ALT (192)
C
967_HG00773_ins
IVNS1ABP normalized total expression
G/G (56)
G/ALT (190)
ALT/ALT (192)
D
967_HG00773_ins
puQTL (prmtr.86427)
Fine-mapped puQTL (prmtr.86427)
eQTL (IVNS1ABP)
-log10 P
PIP
r2
chr1 (Mb)
E
Scale
chr21:
20 kb
hg38
42,460,000
42,470,000
42,480,000
42,490,000
42,500,000
RSPH1
(Ensembl 104 annotation)
RSPH1
(assembled in this study)
52.3098
GM12878 H3K4me3
0.111111
34.4405
GM12878 H3K27ac
0.04
28764_HG02059_ins
(166 bp insertion)
prmtr.70068
prmtr.70069
F
28764_HG02059_ins
RSPH1 normalized promoterr activity
prmtr.70068
prmtr.70069
A/A (47)
A/ALT (208)
ALT/ALT (183)
G
28764_HG02059_ins
RSPH1 normalized total expression
A/A (47)
A/ALT (208)
ALT/ALT (183)
H
28764_HG02059_ins
puQTL (prmtr.70068)
Fine-mapped puQTL (prmtr.70068)
eQTL (RSPH1)
-log10 P
PIP
r2
chr21 (Mb)

**Supplemental Figure 7. Structural variants associated with promoter usage.** (A) The *IVNS1ABP* gene locus. Structures of the *IVNS1ABP* assembled in this study are represented in black with ENCODE GM12878 H3K4me3, H3K4me1, and H3K27ac ChIP-seq signals and H3K27ac HiChIP chromatin interactions. Vertical blue bar indicates the location of an active promoter prmtr.86427. A black bar indicates the location of a structural variant 967_HG00773_ins. (B, C) Comparison of the promoter activities (B) and total expression levels (C) of the *IVNS1ABP* gene among 967_HG00773_ins genotypes. The numbers in parentheses indicate sample size. (D) Associations of puQTL, fine-mapped puQTL for prmtr.86427, and eQTL for *IVNS1ABP* are shown in the top, middle, and bottom panel. 967_HG00773_ins is plotted in a red diamond and colors indicate r-squared values between 967_HG00773_ins and other variants. (E) The *RSPH1* gene locus. Structures of the *RSPH1* in the Ensembl 104 annotation and assembled in this study are represented in black and red, respectively with ENCODE GM12878 H3K4me3 and H3K27ac ChIP-seq signals. Vertical blue bars indicate the location of an active promoters. A black bar indicates the location of a structural variant 28764_HG02059_ins. (F, G) Comparison of the promoter activities (F) and total expression levels (G) of the *RSPH1* gene among 28764_HG02059_ins genotypes. (H) Associations of puQTL, fine-mapped puQTL for prmtr.70068, and eQTL for *RSPH1* are shown in the top, middle, and bottom panel. 28764_HG02059_ins is plotted in a red diamond and colors indicate r-squared values between 28764_HG02059_ins and other variants.

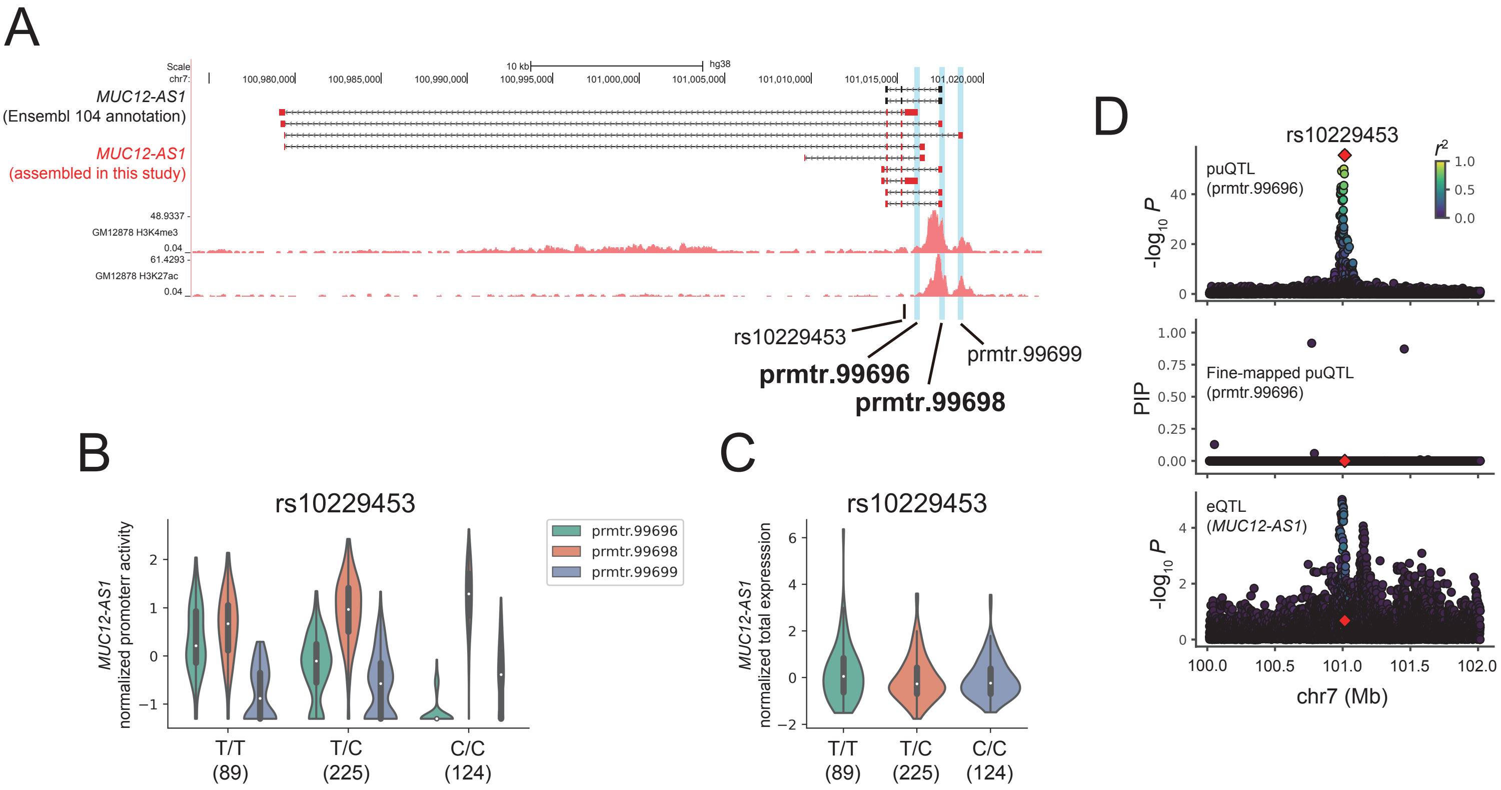


**Supplemental Figure 8. puQTL associated with distinct promoters of the *MUC12-AS1* gene with opposite effects.** (A) The *MUC12-AS1* gene locus. Structures of the *MUC12-AS1* in the Ensembl 104 annotation and assembled in this study are represented in black and red, respectively, with ENCODE GM12878 H3K4me3 and H3K27ac ChIP-seq signals. Vertical blue bars indicate the location of active promoters. A black bar indicates the location of a variant rs10229453. (B, C) Comparison of the promoter activities (B) and total expression levels (C) of the *MUC12-AS1* gene among rs10229453 genotypes. The numbers in parentheses indicate sample size. (D) Associations of puQTL, fine-mapped puQTL for prmtr.99696, and eQTL for *MUC12-AS1* are shown in the top, middle, and bottom panel. rs10229453 is plotted in a red diamond and colors indicate r-squared values between rs10229453 and other variants.

# A

```
CLUSTAL multiple sequence alignment by MUSCLE (3.8)

sp|Q9BPZ7-2|SIN1_HUMAN      MAFLDNPTIILAHIRQSHVTSDDTGMCEMVLIDHDVDLEKIHPPSMPGDSGSEIQGSNGE
tr|B1AMA5|B1AMA5_HUMAN      ------------------------------------------------------------

sp|Q9BPZ7-2|SIN1_HUMAN      TQGYVYAQSVDITSSWDFGIRRRSNTAQRLERLRKERQNQIKCKNIQWKERNSKQSAQEL
tr|B1AMA5|B1AMA5_HUMAN      ------------------------------------------------------------

sp|Q9BPZ7-2|SIN1_HUMAN      KSLFEKKSLKEKPPISGKQSILSVRLEQCPLQLNNPFNEYSKFDGKGHVGTTATKKIDVY
tr|B1AMA5|B1AMA5_HUMAN      ------------------------------------------------------------

sp|Q9BPZ7-2|SIN1_HUMAN      LPLHSSQDRLLPMTVVTMASARVQDLIGLICWQYTSEGREPKLNDNVSAYCLHIAEDDGE
tr|B1AMA5|B1AMA5_HUMAN      ------------MTVVTMASARVQDLIGLICWQYTSEGREPKLNDNVSAYCLHIAEDDGE
                                        ************************************************

sp|Q9BPZ7-2|SIN1_HUMAN      VDTDFPPLDSNEPIHKFGFSTLALVEKYSSPGLTSKESLFVRINAAHGFSLIQVDNTKVT
tr|B1AMA5|B1AMA5_HUMAN      VDTDFPPLDSNEPIHKFGFSTLALVEKYSSPGLTSKESLFVRINAAHGFSLIQVDNTKVT
                            ************************************************************

sp|Q9BPZ7-2|SIN1_HUMAN      MKEILLKAVKRRKGSQKVSGSRADGVFEEDSQIDIATVQDMLSSHHYKSFKVSMIHRLRF
tr|B1AMA5|B1AMA5_HUMAN      MKEILLKAVKRRKGSQKVSGSRADGVFEEDSQIDIATVQDMLSSHHYKSFKVSMIHRLRF
                            ************************************************************

sp|Q9BPZ7-2|SIN1_HUMAN      TTDVQLGISGDKVEIDPVTNQKASTKFWIKQKPISIDSDLLCACDLAEEKSPSHAIFKLT
tr|B1AMA5|B1AMA5_HUMAN      TTDVQLGISGDKVEIDPVTNQKASTKFWIKQKPISIDSDLLCACDLAEEKSPSHAIFKLT
                            ************************************************************

sp|Q9BPZ7-2|SIN1_HUMAN      YLSNHDYKHLYFESDAATVNEIVLKVNYILESRASTARADYFAQKQRKLNRRTSFSFQKE
tr|B1AMA5|B1AMA5_HUMAN      YLSNHDYKHLYFESDAATVNEIVLK-----------------------------------
                            *************************

sp|Q9BPZ7-2|SIN1_HUMAN      KKSGQQ
tr|B1AMA5|B1AMA5_HUMAN      ------
```

: N-terminal domain

: Conserved region in middle domain

: Preckstrin-homology domain

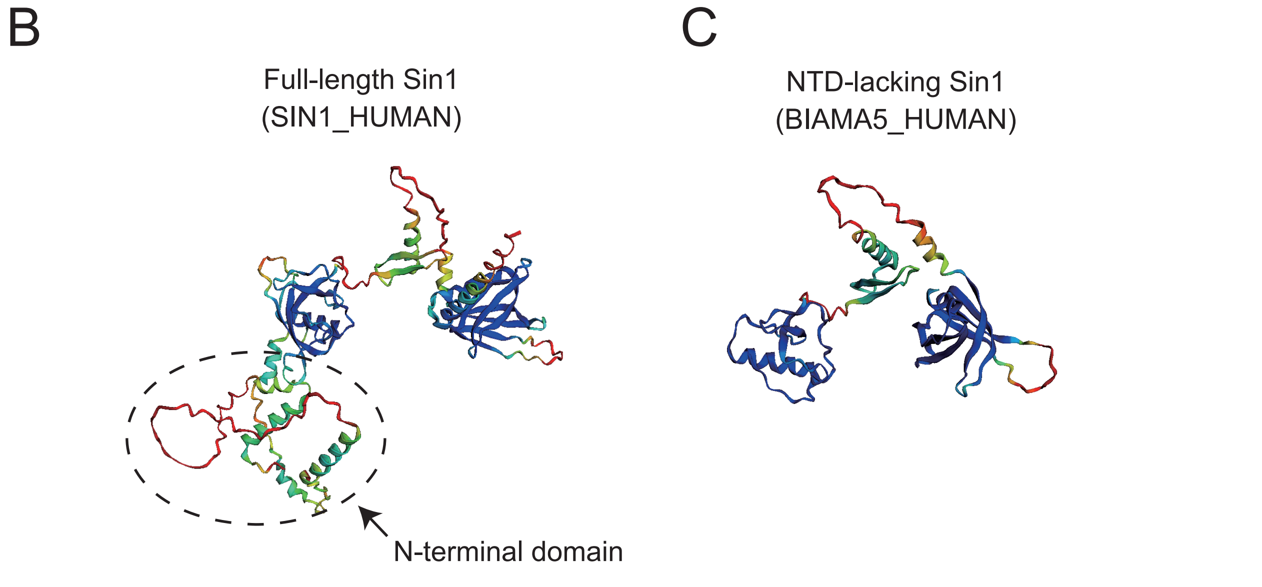


**Supplemental Figure 9. Comparison of the Sin1 proteins translated from distinct transcripts.** (A) Multiple sequence alignment of two isoforms of the Sin1 (MAPKAP1) protein. Protein sequences of full-length Sin1 (SIN1_HUMAN) and NTD-lacking Sin1 (BIAMA5_HUMAN) are alingned in top and bottom rows, respectively. Asterisks (*) indicate positions of matched residues. Red, black, and blue rectangles indicate sequences of N-terminal domain, conserved region in middle domain, and Preckstrin-homology domain, respectively. (B, C) Protein structures of full-length Sin1 (SIN1_HUMAN) (B) and NTD-lacking Sin1 (BIAMA5_HUMAN) (C) predicted by AlphaFold2 with colors representing per-residue confidence score. Dotted circles indicate N-terminal domain.
